# Top predator introduction changes the effects of spatial isolation on freshwater community structure

**DOI:** 10.1101/857318

**Authors:** Rodolfo Mei Pelinson, Mathew A. Leibold, Luis Schiesari

## Abstract

The importance of local selective pressures on community structure is predicted to increase with spatial isolation when species favored by local conditions also have higher dispersal rates. In freshwater habitats, the introduction of predatory fish can produce trophic cascades because fish tend to prey upon intermediate predatory taxa, such as predatory insects, which indirectly benefits herbivores and detritivores. Similarly, spatial isolation is known to limit predatory insect's colonization rates more strongly than of herbivores and detritivores, thus generating similar effects. Here we tested the hypothesis that the effect of introduced predatory fish on macroinvertebrate community structure increases across a gradient of spatial isolation by conducting a field experiment where artificial ponds with and without fish (the Redbreast Tilapia) were constructed at three different distances from a source wetland. Overall results show that fish do reduce the abundance of predatory insects but have no effect on the abundance of herbivores and detritivores. Spatial isolation, however, does strengthen the trophic cascade caused by dispersal limitation of predatory insects, but only in the absence of fish. More importantly, macroinvertebrate communities with and without fish tend to diverge more strongly at higher spatial isolation, however, this pattern was not due to an increase in the magnitude of the effect of fish, as initially hypothesized, but to a change in the effect of isolation in the presence of fish. We argue that as spatial isolation increases, suitable prey, such as predatory insects become scarce and fish increases predation pressure upon herbivores and detritivores, dampening the positive effect of spatial isolation on them. Our results highlight the importance of considering interspecific variation in dispersal and multiple trophic levels to better understand the processes generating metacommunity structures.

## Introduction

The presence of top predators has been widely recognized to cause strong changes in species abundance patterns and community structure (*e.g.* Paine 1966, Holt et al. 1994, Leibold 1996, Frank et al. 2005, Newsome et al. 2017). Less frequently acknowledged, however, is that such changes, as of any other local selective pressure, can be modulated by dispersal (Leibold and Chase 2018, Guzman et al. 2019). Classic metacommunity theory recognizes that the frequency and intensity of dispersal can modulate the relative importance of ecological drift (*i.e.* demographic stochasticity) and niche selection processes, such as the presence of top predators, in structuring metacommunities (Leibold et al. 2004). When species dispersal rates are low for all species involved, or patches are isolated, stochastic events are likely to cause communities to drift towards multiple different compositions that aren’t necessarily related to these local selective pressures (Leibold and Chase 2018). In contrast, if species dispersal rates are sufficiently high for all species involved or patches are highly connected, the constant arrival of migrants should override the effects of drift and niche selection making communities more similar to each other (*i.e.* the mass-effect; Mouquet and Loreau 2003, Leibold and Chase 2018). Thus, the presence of a top predator would be expected to cause greater change in prey community structure at intermediate levels of connectivity. However, these predictions may not be realized if there is interspecific variation in dispersal rates (Vellend et al. 2014, Guzman et al. 2019). Variation in dispersal can actually intensify the consequences of niche selection if the traits that confer higher fitness within a set of local environmental conditions (*e.g.* ability to escape predation) are positively correlated with traits that confer higher dispersal rates (Vellend et al. 2014).

Freshwater communities are strongly influenced by the presence of predatory fish (Hrbáček 1962, Brooks and Dodson 1965, Wellborn et al. 1996). In the absence of fish, predatory invertebrates such as aquatic beetles and dragonfly larvae are often top predators. Compared to fish, they are usually less efficient, gape-limited sit-and-wait predators that consume smaller prey (Wellborn et al. 1996). However, when present, fish, which are generally large visually-oriented predators, tend to preferentially consume large prey, which frequently happen to be those predatory insects (Wellborn et al. 1996, McCauley 2008). Therefore, by preferentially consuming predatory insects, fish can produce a trophic cascade that results in an increase in the abundance of small herbivores and detritivores (Diehl 1992, Goyke and Hershey 1992).

Freshwater insects can also vary substantially in dispersal and colonization rates and such variation is often correlated with trophic level (Bilton et al. 2001, Shulman and Chase 2007, Chase and Shulman 2009, Shurin et al. 2009, Guzman et al. 2019). For example, predatory insects tend to have larger body sizes than their prey, and thus higher locomotory ability (McCann et al. 2005). However, predatory insects also tend to have smaller population sizes (Cohen et al. 2003) and longer generation times than their prey, which makes colonization events in spatially isolated ponds rarer (Chase and Shulman 2009). They can also be indirectly disfavored by habitat isolation if their prey is dispersal-limited or unable to reach high population sizes (Hein and Gillooly 2011). Insects that are herbivores and detritivores, by contrast, tend to have shorter generation times and larger population sizes, which can increase their number of dispersal events when compared to predatory insects (Poff et al. 2006). Moreover, the smaller body sizes of herbivores and detritivores may greatly expand their dispersal range by wind transport (Muehlbauer et al. 2014). Therefore, the relative decrease in colonization rates across a gradient of spatial isolation may be stronger for predatory insects. An important outcome of this steeper decline in colonization rates for higher trophic level insects is that spatial isolation can promote changes in community structure that are essentially analogous to trophic cascades, as the abundance of predators would decline and that of herbivores and detritivores increase in more isolated habitats (Shulman and Chase 2007, Chase and Shulman 2009). Therefore, because traits that confer lower dispersal rates to predatory insects are correlated with traits that make them susceptible to predation by fish, spatial isolation is likely to strengthen the cascading effect caused by the presence of fish.

Our study aimed at experimentally assessing whether and how spatial isolation can change the effects of the introduction of a generalized fish predator on freshwater insect community structure. We designed our study to test the following hypotheses with special attention to the role of interspecific variation in dispersal: First, the presence of predatory fish would change insect communities mostly as a consequence of a cascading effect by preferentially preying upon larger predatory insects, decreasing their abundance (Fig. 1a) and causing the abundance of herbivores and detritivores to increase (Fig. 1b). Second, spatial isolation would promote a change in community structure due to a similar cascading effect, by reducing the abundance of predatory insects (Fig. 1a), which can successfully reach more isolated habitats less frequently than their prey, also increasing the abundance of herbivores and detritivores (Fig. 1b). Finally, and third, increasing isolation should intensify the effect of fish on community structure because the ecological traits that strengthen the cascading effects in both cases are considered positively correlated (Fig. 1c).

**Figure 1.**
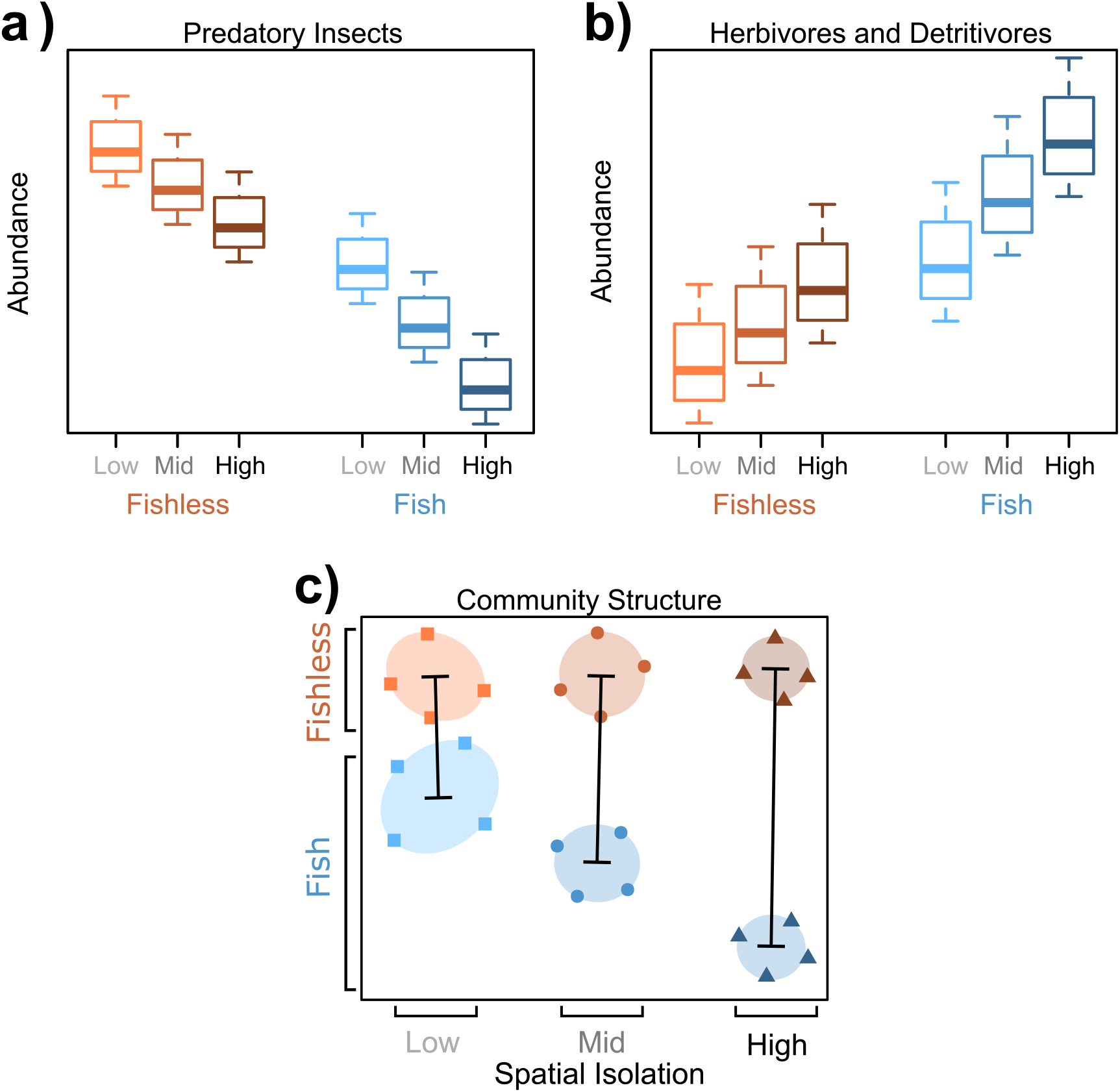
Graphical representation describing our initial hypotheses. A: Both fish and spatial isolation are predicted to have negative effects on the abundance of predatory insects. This is expected because fish are visually oriented predators, thus they should preferentially prey upon large-bodied predatory insects. Similarly, even though predatory insects often have higher dispersal ability, their dispersal rates are lower than of herbivores and detritivores because they have lower population sizes, thus fewer dispersal events. B: In contrast and as a consequence of the reduction in the abundance of predatory insects, the abundance of herbivores and detritivores are predicted to increase in the presence of fish and as spatial isolation increases. C: If species vary in dispersal rates and good dispersers are favored by local conditions, as shown in B, or bad dispersers are disfavored by local conditions, as shown in A, the importance of niche selection processes (*e.g.* the effect of fish) may increase with spatial isolation, increasing the differences between communities with and without fish (black bars). C is a hypothetical ordination plot where each symbol represents a community and similar symbols are communities from the same treatment.

## Methods

We conducted our field experiment at the Estação Ecológica de Santa Bárbara (EESB) in Águas de Santa Bárbara, São Paulo, Brazil (22°48’59” S, 49°14’12” W). The EESB is a 2,712-ha protected area predominantly covered with open savanna habitats, with smaller portions of seasonal semideciduous forests, *Pinus* spp., and *Eucalyptus* spp. plantations (Melo and Durigan 2011). Soils are sandy and climate is Koeppen’s Cwa, i.e., warm temperate with dry winters and hot summers (Kottek et al. 2006). Mean annual rainfall is ~1250mm with a distinct rainy season from October to March (January being the wettest month with ~200mm rainfall) and a dry season from April to September (July being the driest month with ~25mm rainfall; (Hijmans et al. 2005). In the EESB the experiment was implemented in an area covered by second-growth cerrado *sensu stricto*, a moderately dense, open-canopy savanna habitat (Melo and Durigan 2011).

Experimental units consisted of ~1,200L artificial ponds dug into the ground and lined with a 0.5 mm thick, high-density polyethylene geomembrane to retain water. Each pond was 4 m long, 1m wide and 40 cm deep. Walls were vertical along the length of the pond; 1 m-long ramps terminating at ground level at each short side of the pond provided shallow microhabitats for freshwater organisms and escape for terrestrial fauna that eventually fell into the water. Two roof tiles were placed at the waterline in each of the short sides to provide shelter and/or oviposition habitat. Three 30 cm-long, 10 cm-wide PVC pipes were placed in the water to provide shelter for fishes (see Appendix S1).

### Experimental design

The experiment followed a fully factorial design crossing fish presence (presence/absence) with spatial isolation (three levels of isolation). The isolation treatment was achieved by establishing 8 artificial ponds along each of three parallel transects 30m, 120m, and 480m from a source wetland consisting of a stream (Riacho Passarinho) and its marshy floodplain (see Appendix S1). Within each transect, the distance between adjacent artificial ponds was 30 m. The artificial ponds were naturally colonized by organisms dispersing in the landscape including phytoplankton, zooplankton, and macroinvertebrates. The chosen distances from the source habitat and their spacing intervals represent a steep spatial isolation gradient for most aquatic insects (Wilcox 2001, Trekels et al. 2011; see Appendix S2). The well-drained sandy soils ensured that no other ponds and puddles formed during the rainy season at our study site, making the near stream and its floodplain the most likely source of aquatic insects colonizing the artificial ponds. Each fish-by-distance treatment was replicated four times for a total of 24 artificial ponds.

The experiment ran from 18-Jan-2017 to 24-Apr-2017. Between 18 and 25-Jan-2017 mesocosms were filled with well water, which does not contain already established plankton or macroinvertebrate communities. On 28-Jan-2017 we added to each mesocosm 1000g (wet mass) of leaf litter composed of equal amounts of grass and tree leaf litter to provide structural complexity for benthic organisms. On 29-Jan-2017 we added to each mesocosm 15g of dog chow to provide an initial pulse of nutrients. At the same day we added one Redbreast Tilapia (*Coptodon rendalli*, standard length 99.2 mm ± 5.9 mm, wet mass 40.2 g ± 8.8 g, mean ± SD, N=12) per fish treatment pond, collected in a small reservoir outside the EESB. Over the course of the experiment, we monitored ponds for fish presence, replacing missing fish as soon as noticed. Fish dying within the first days of the experiment, possibly due to handling stress, were immediately replaced. In the following weeks, mesocosms water became turbid and it was not always possible to assess fish presence without netting. Because netting could represent a considerable disturbance to freshwater communities, we waited until the end of each sampling survey to seine the ponds and thereby assess fish presence. Two fishes were found to be missing by the end of the first and second surveys (see Appendix S3). No fish died between the second and third surveys. Therefore, every pond with fish had at least 4 weeks of fish presence prior to the final survey.

We selected the Redbreast Tilapia for three reasons. First, tilapias are generalized omnivorous predators that, even though many times are described as having a preference for feeding upon macrophytes (Rao et al. 2015) and zooplankton (Lazzaro 1991), can also prey upon macroinvertebrates when available (Chimbari et al. 1997; also confirmed in a pilot laboratory experiment we conducted, see Appendix S4). Second, they are hardy species capable of surviving in a wide range of environmental conditions including low oxygen levels and a broad range of temperatures (Caulton 1977, Tran-Duy et al. 2008), conditions likely to be found in our shallow artificial ponds. Third, we understand that the spatial context at which natural and constructed ponds occur is important when assessing the impacts of invasive Tilapias on freshwater communities. The Redbreast Tilapia is, along with the Nile Tilapia (*Oreochromis niloticus*), one of the most widely introduced fishes in the world for aquaculture and recreational fisheries (Britton and Orsi 2012). These African species represented ~11% (6.1 million tons) of the entire freshwater fish production in the world and ~40% (0.6 million tons) in the Americas in 2017 (FAO 2019). In Brazil, Redbreast and Nile Tilapias are found in reservoirs and lakes in most river basins, and their spread to new river basins may be a matter of time considering that their stocking is still encouraged by public policies (Zambrano et al. 2011, Britton and Orsi 2012, Pelicice et al. 2014, Daga et al. 2016). Indeed, a very common land management practice in rural Brazil is the construction of dugout or impounded lakes, where the Tilapia is usually the first choice of fish species for stocking.

### Freshwater community sampling surveys

To assess the influence of fish presence, spatial isolation, and their interaction on community assembly we conducted three sampling surveys of the artificial ponds after ~3 weeks (18 to 23-Feb-2017), ~8 weeks (23 to 27-Mar-2017), and ~12 weeks (20 to 24-Apr-2017) of the experiment. Freshwater communities were dominated by insects, which were sampled by sweeping half of the pond twice, including both pelagic and benthic habitats, with a hand net (mesh size 1.5 mm). Samples were cleaned of debris and stored in 70% ethanol. We identified and counted all aquatic macroinvertebrates to the lowest reliable taxonomical level using taxonomic keys for South American freshwater insects (Costa et al. 2004, Pereira et al. 2007, Segura et al. 2011, Hamada et al. 2014).

### Data analysis

We tested whether the presence of fish, spatial isolation, sampling survey, and their interaction caused any changes in total abundance of predatory insects, and herbivores and detritivores separately through sequential likelihood ratio tests. We modeled species total abundance using generalized linear mixed models (*i.e.* GLMMs) with a negative binomial distribution, and pond identifications as a random effect term. We also performed pairwise *post-hoc* tests with Šidák adjustment of p-values for factors with more than two levels, and for interactions, when they were important in explaining abundance patterns.

Prior to the analysis of community structure, we removed species that occurred in less than two ponds considering each sampling survey (*i.e.* singletons and doubletons) both because they are uninformative to general community patterns and because they complicate model parameter estimation (Warton et al. 2015a). To test the hypothesis that community structure is influenced by the presence of fish, spatial isolation, and their interaction, we performed sequential likelihood ratio tests on multivariate community data (Warton et al. 2015b) assuming that species abundances come from a negative binomial distribution due to observed overdispersion in the data. The main advantage of this approach, in opposition to a perMANOVA, is the possibility of accounting for the mean-variance relationship of abundance data, and the better interpretability of change on community structure by looking at coefficients of the effect of treatments on individual species and their respective confidence intervals. Additionally, we can test for the effect of traits on species responses to treatments (*i.e.* model-based fourth corner solution; Brown et al. 2014, Warton et al. 2015a). Thus, we also tested whether trophic level (*i.e.* strict predator VS herbivores and detritivores) is a good predictor of the changes in community structure through likelihood ratio tests. Note here that this analysis differs from tests of total abundance because it tests for similar responses to treatments of species in each trophic level, whereas total abundances tend to show patterns that are driven by the more abundant species. A shortcoming of this approach lies in the difficulties of modeling correlations in species abundances (Warton et al. 2015a, 2015b). To account for those correlations when computing p-values for the entire communities, we performed 10,000 bootstraps resamples, using the PIT-trap bootstrap resample procedure (Warton et al. 2017), shuffling entire rows of the incidence matrix (ponds), keeping species abundances in the same ponds always together. To account for lack of independence between the same ponds sampled across time when analyzing all surveys together, ponds were considered blocks, so in each permutation step we shuffled ponds freely within blocks (*i.e.* only across time), then we shuffled the entire bocks freely (*i.e.* across fish and isolation treatments). Because we found significantly different effects of fish and isolation treatments across different sampling surveys, we repeated the analysis in each sampling surveys separately.

Due to inevitable practical constraints, one limitation of our experimental design is that the experimental units were not fully spatially isolated from each other within the isolation treatments, meaning that they were not fully independent from each other. Therefore, we also tested for the presence of spatial dependence among experimental units using Moran’s Eigenvector Maps (MEMs) (Borcard and Legendre 2002, Dray et al. 2006). We did not find important effects of neighbor communities on community structures (see Appendix S5 for details).

In the community structure analysis, a significant interaction between fish and isolation means that there is either or both a difference in direction (*i.e.* positive or negative effect) or magnitude of the effect of fish in different isolation treatments. To specifically test for differences in the size of the effect of fish, regardless of direction, we performed a model based unconstrained ordination via generalized linear latent variable models (GLLVM; Niku et al. 2017) with a negative binomial distribution using two latent variables for each of the sampling surveys (Hui et al. 2015). The latent variables were estimated via variational approximation (Hui et al. 2017). Here the latent variables are interpreted as the axis of any ordination analysis, such as non-metric multidimensional scaling (*i.e.* nMDS) (Hui et al. 2015). After performing the ordination, we computed the centroids of each treatment group and the distance between the centroids of fish and fishless treatments in each isolation treatment as a measure of the size of the effect of fish (black bars in Fig. 3). Then we tested whether this distance is significantly different across all the isolation treatments. To test for that we designed a permutation procedure to only permute ponds across isolation treatments, keeping the fish treatment constant. This represented a null scenario where the effect of fish is the same in all isolation treatments. For this analysis, we corrected p-values for multiple comparisons using the false discovery rate correction (FDR). We also used those ordinations to visualize the effect of treatments on community structure.

We had to exclude samples from four ponds in the third survey due to mislabeling, therefore we analyzed 24 ponds for the first and second surveys and 20 for the third one. In the last survey treatments with fish in 30 m, 120 m, and 480 m, and without fish in 480 m, had only three replicates (see Appendix S4). All analyses were implemented in software R version 4.0.2. (R Core Team 2020). Code and data to reproduce the analyses and figures are available as an R package on github in the following repository: RodolfoPelinson/PredatorIsolationComm.

## Results

### Total Abundance

Detailed information about the taxa sampled across the experiment is available in Appendix S6. The total abundance of predatory insects increased over the course of the experiment, decreased in the presence of fish, and decreased with spatial isolation (Table 1; Fig. 2a). The total abundance of herbivores and detritivores also increased throughout the experiment, but fish did not affect it. Spatial isolation, however, did increase the total abundance of herbivores and detritivores, but only in the last survey. This effect was stronger for fishless ponds (Table 1; Fig. 2b).

**Table 1.**
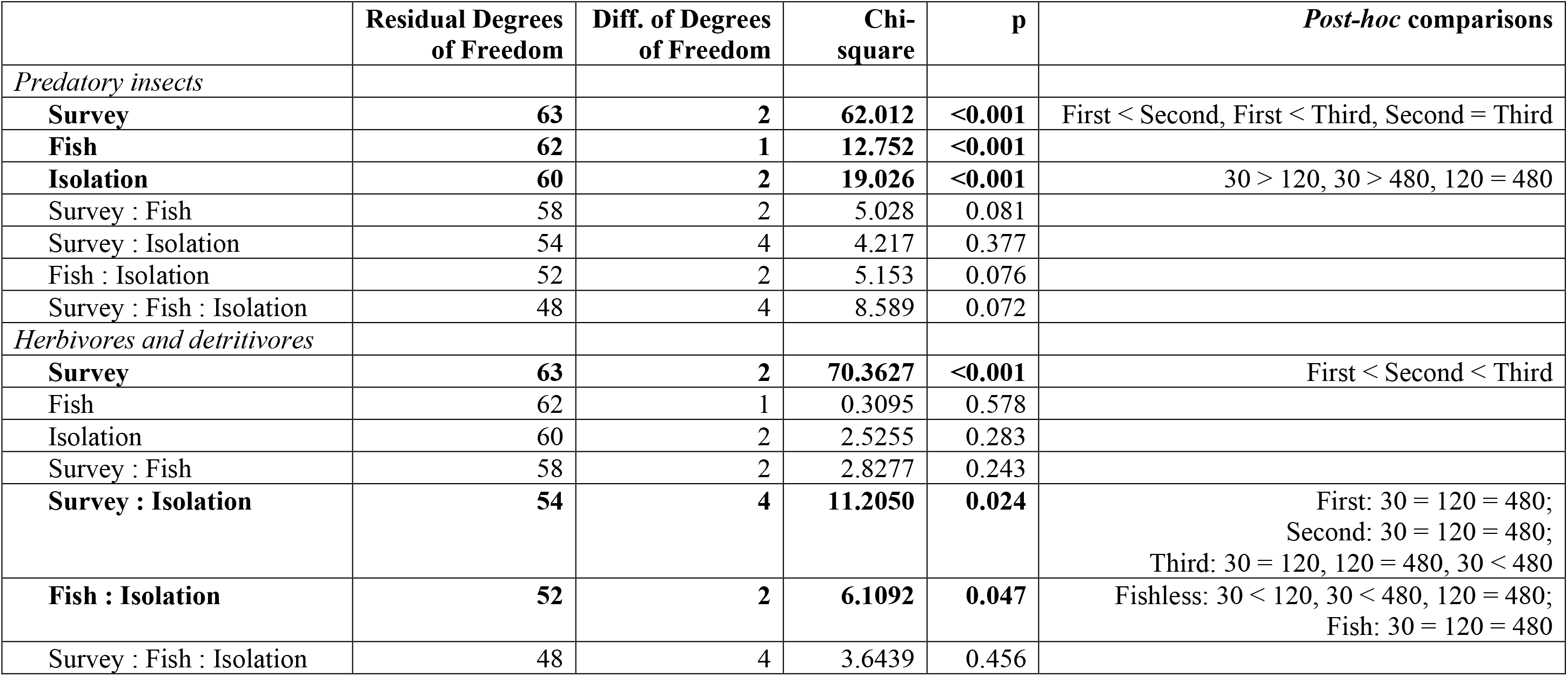
Summary of likelihood ratio tests for total abundances of predatory insects and herbivores and detritivores. Bold values represent a significant effect (p < 0.05).

**Figure 2.**
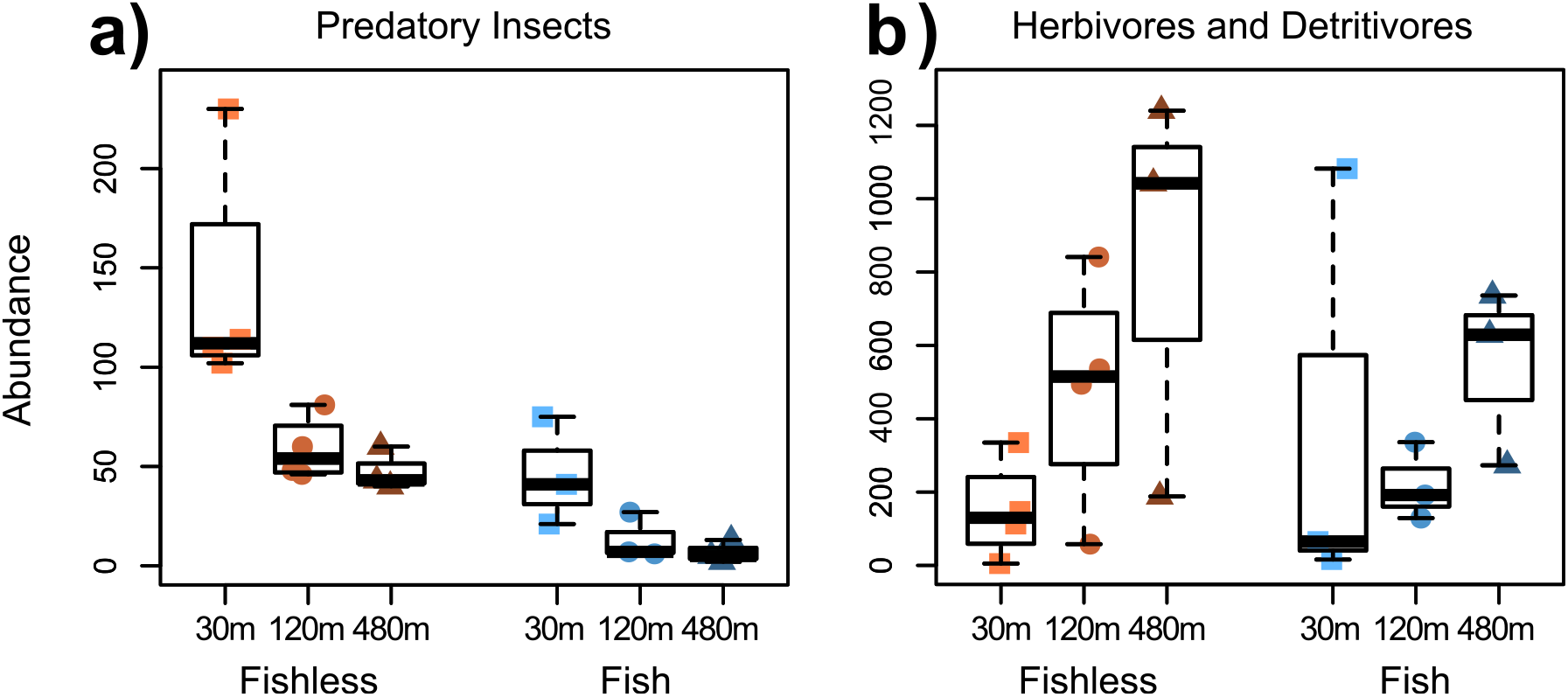
Total abundance of predatory insects (A) and herbivores and detritivores (B) for the last survey. Plots for the first and second surveys are provided in Appendix S7.

### Community Structure

The effect of treatments on community structure became clearer and stronger over time (Table 2). We, therefore, will focus on the results of the last (third) survey for clarity. Community structure was generally affected by the interaction of the presence of fish and spatial isolation. More importantly, this interactive effect of treatments was significantly explained by trophic level (Table 2; Fig. 3). Predatory insects had similar negative responses to the presence of fish, but only in 120 and 480 m (Fig. 4a). The general lack of this effect at 30 m was mostly because *Pantala* dragonflies suffered no effect of fish and *Orthemis* dragonflies were positively affected by it at this distance (Fig. 3; see Appendix S9). Predatory insects also had coherent negative responses to isolation from 30 to 120 and 480 m distances, but only in ponds with fish (Fig. 4b and 4c). Different from observed for total abundance, in fishless ponds, some predatory insects, such as *Pantala* and *Orthemis* dragonflies, were positively affected (Fig. 3; see Appendix S9). Herbivores and detritivores only responded as a group to the presence of fish at the intermediate spatial isolation, and this response, contrary to what was expected, was negative (Fig. 4a). However, they were positively affected by isolation from 30 m to 120 m and 480 m, but only in fishless ponds (Fig. 4c).

**Figure 3.**
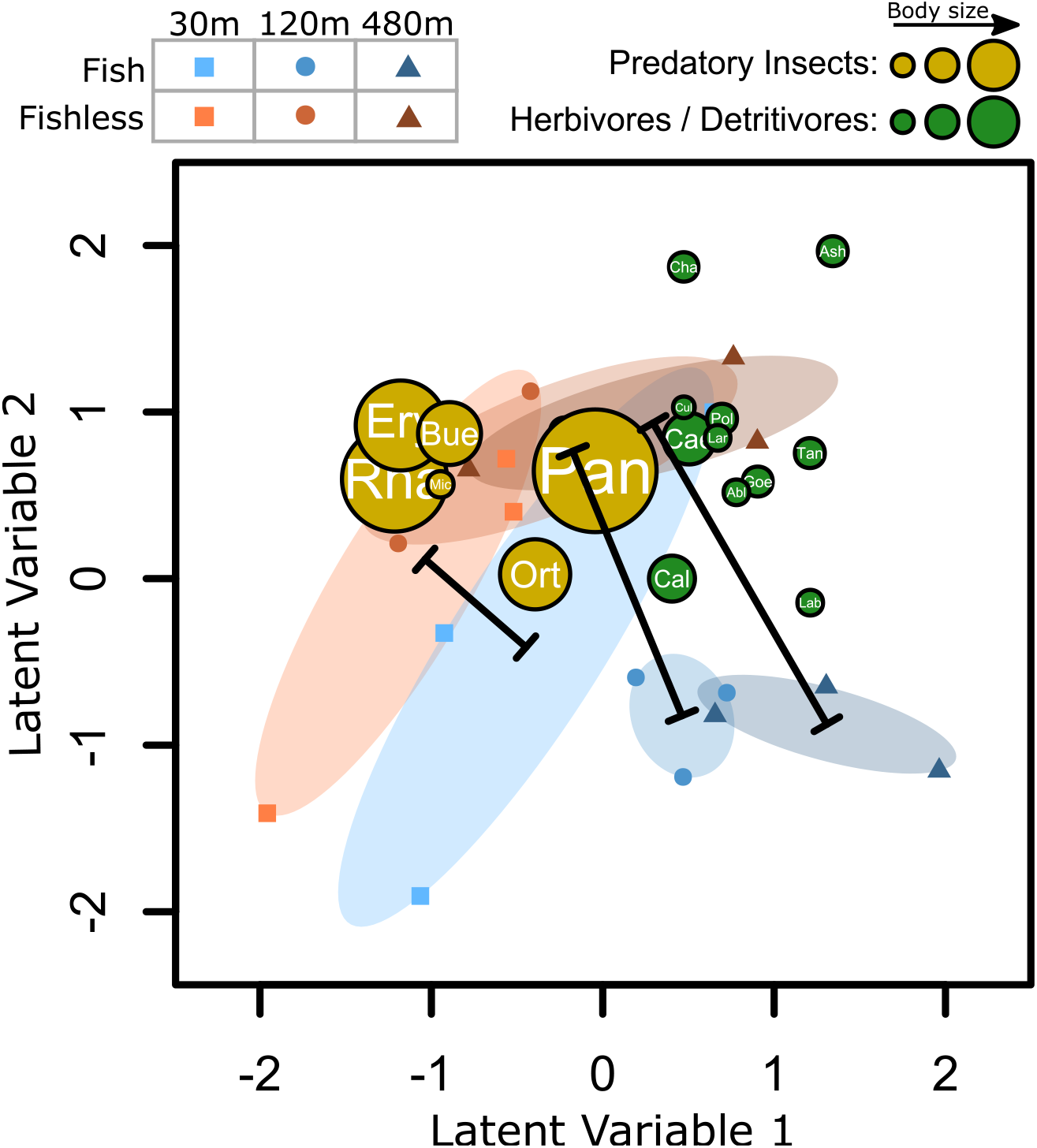
Unconstrained ordination showing pond communities (symbols) and species (filled circles) for the third survey. Yellow filled circles represent predatory insects and green filled circles represent herbivores and detritivores. The sizes of the circles are proportional to maximum body size (the volume of the largest individual of each species in a log-scale). Abbreviations of names of taxa are provided in Appendix S6. Symbols close to each other have more similar community structures than those that are further apart. Species are positioned closer to treatments where their abundance is higher. Note that herbivores and detritivores are more strongly associated with high isolation treatments, especially fishless ponds, whereas predatory insects are associated with low isolation treatments, especially fishless ponds. Note also that *Pantala* and *Orthemis* dragonflies (Pan and Ort) are detached from the other predatory insects, *Pantala* being closer to highly isolated fishless treatments and *Orthemis* to ponds with fish in low isolation. Black bars show the distance between fish and fishless treatment’s centroids in each isolation treatment. Similar ordination plots for the first and second surveys are provided in Appendix S8.

**Table 2.** Summary of likelihood ratio tests for community structure considering all sampling surveys together and separately. Bold values represent a significant effect (p < 0.05).

|  | **Residual**<br>**Degrees of**<br>**Freedom** | **Diff. of**<br>**Degrees of**<br>**Freedom** | **Deviance** | **p** |
| --- | --- | --- | --- | --- |
| *All Sampling Surveys* |  |  |  |  |
| **Survey** | **57** | **2** | **392.8** | **<0.001** |
| **Fish** | **56** | **1** | **89.2** | **<0.001** |
| **Isolation** | **54** | **2** | **109.2** | **0.001** |
| **Fish : Isolation** | **52** | **2** | **120.3** | **<0.001** |
| Survey : Fish | 50 | 2 | 50.9 | 0.075 |
| Survey : Isolation | 46 | 4 | 78.5 | 0.247 |
| **Survey : Fish : Isolation** | **42** | **4** | **81.2** | **0.014** |
| *1st Sampling Survey* |  |  |  |  |
| Fish | 22 | 1 | 19.01 | 0.104 |
| Isolation | 20 | 2 | 24.38 | 0.398 |
| **Fish : Isolation** | **18** | **2** | **42.60** | **0.024** |
| (Fish : Isolation) :Trophic Level | 197 | 5 | 19.26 | 0.106 |
| *2nd Sampling Survey* |  |  |  |  |
| **Fish** | **22** | **1** | **62.28** | **0.002** |
| **Isolation** | **20** | **2** | **71.81** | **0.025** |
| **Fish : Isolation** | **18** | **2** | **72.15** | **0.016** |
| **(Fish : Isolation) :Trophic Level** | **404** | **5** | **33.74** | **0.002** |
| *3rd Sampling Survey* |  |  |  |  |
| **Fish** | **18** | **1** | **49.09** | **0.018** |
| Isolation | 16 | 2 | 72.96 | 0.055 |
| **Fish : Isolation** | **14** | **2** | **91.12** | **0.015** |
| **(Fish : Isolation) :Trophic Level** | **332** | **5** | **33.71** | **0.028** |

**Figure 4.**
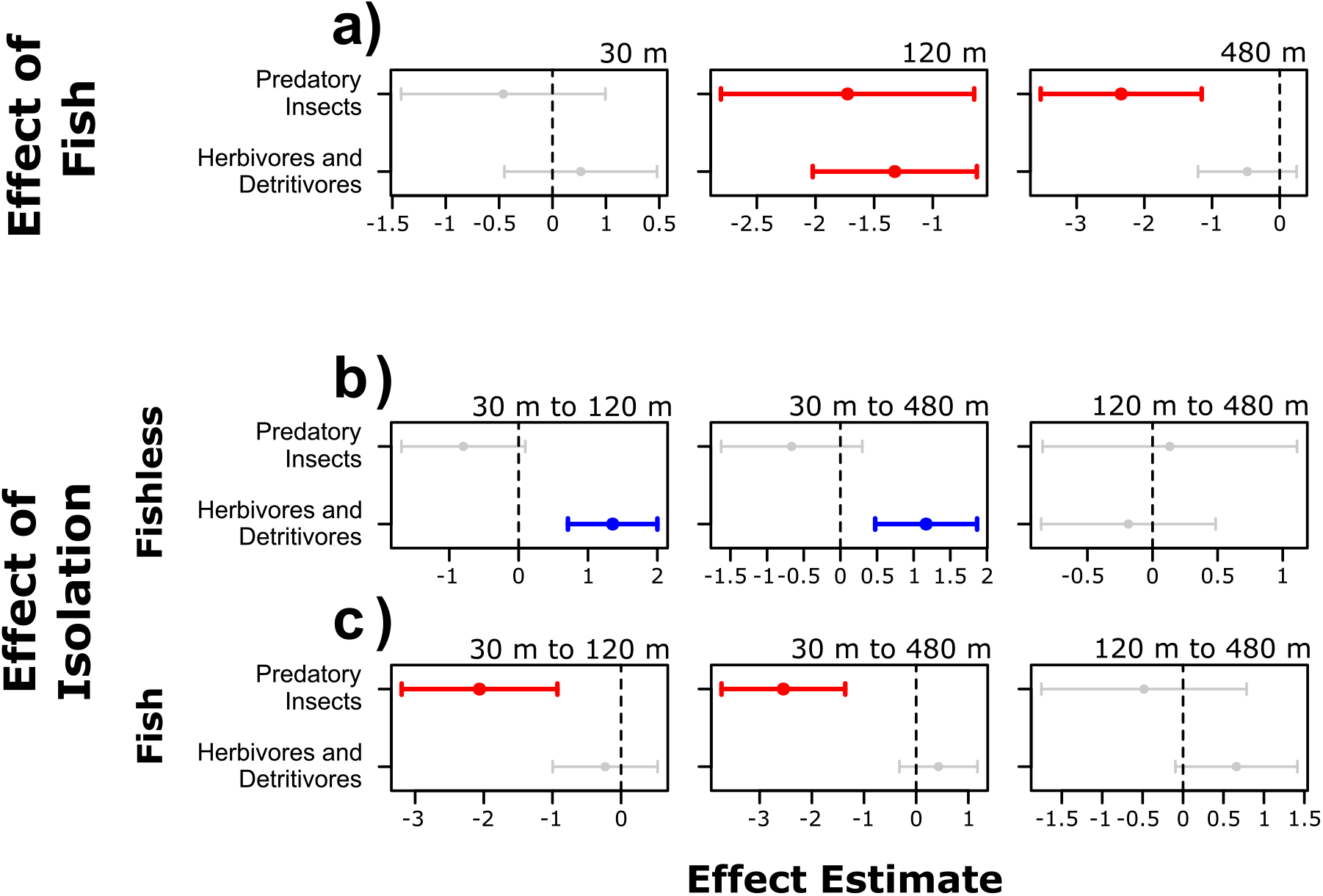
95% Confidence intervals for the effect of fish and isolation on the abundance of predatory insects and herbivores and detritivores when comparing pairs of treatments for the third survey of the experiment. Confidence intervals not crossing the zero hatched line were considered significant effects and colored; blue bars represent positive and red bars negative responses to a given treatment when compared to a reference treatment. A are effects of the presence of fish in each isolation treatment. B are effects of isolation in fishless ponds and C in ponds with fish. In each of the B and C panels, we show the effects of increasing isolation from 30 to 120 m, from 30 to 480 m, and from 120 m to 480 m

When we tested for differences in the distance between centroids of fish and fishless communities for the third survey, we found that it was larger at 480 m than at 30 m (Difference in distance between centroids: 1.63; adj. p: 0.038; Fig. 3) meaning that, as we expected, fish and fishless ponds are more different from each other in more isolated ponds. The distance between centroids at 120 m was not significantly different from 30 m (Difference in distance between centroids: 1.25; adj. p: 0.098; Fig. 3) or 480 m (Difference in distance between centroids: 0.38; adj. p: 0.601; Fig. 3).

## Discussion

Generally, both the presence of fish and spatial isolation had important effects on freshwater community structure and responses to treatments were different for different trophic levels. More importantly, our results, as expected, diverged from classic metacommunity models, which predicts that the effect of local niche selection on community structure should be stronger at intermediate levels of isolation (Mouquet and Loreau 2003, Leibold et al. 2004, Leibold and Chase 2018). However, even though we show that the difference between communities with and without fish only increases with isolation, it does not happen because of a positive correlation in the responses of predatory insects and herbivores and detritivores to the presence of fish and isolation.

The presence of fish indeed shifted species composition through a reduction of predatory insects, notably dytiscid beetles, notonectids, and dragonflies, all relatively large taxa (see Appendix S9). Contrary to our expectations and previous work (*e.g.* Goyke and Hershey 1992), however, it did not translate into an increase in abundance of herbivores and detritivores. Previous work has shown that even though generalist predatory fish frequently prey upon multiple trophic levels, its effect on predatory insects tend to be disproportionally larger, as they are easier to catch and have smaller population sizes (Diehl 1992, Batzer et al. 2000). Thus, the net effect of fish on the abundance of herbivores and detritivores is a sum of both direct negative and indirect positive effects (Batzer et al. 2000). In our experiment, the direct negative effect of fish predation upon herbivores and detritivores might have balanced out the indirect positive effect of the reduced abundance of predatory insects.

We initially hypothesized that the abundance of predatory insects should decrease as spatial isolation increases, and the abundance of their prey should increase in response. We partially found support for this hypothesis. The total abundance of predatory insects does decrease as spatial isolation increases, however, not all predatory insects appear to be dispersal limited. Predatory insects taxa only consistently negatively responded to isolation when fish was present. When it was absent, there were two clear exceptions to this pattern: *Pantala* and *Orthemis* dragonflies. While other predators, such as diving beetles (*i.e. Rhantus*) and water striders (*i.e. Microvelia*), exhibited a strong decrease in abundance in both fish and fishless ponds as spatial isolation increased, both of these dragonflies were positively affected by isolation in the absence of fish. Dragonflies are known to have a great dispersal ability; thus it is not surprising that at the scale of our experiment they would suffer small or no negative effects at all of isolation (McCauley 2006). We believe that the positive effects observed might be a net consequence of the absence of other predatory insects in more isolated ponds, which could have released those dragonflies from competition or made these ponds more attractive for adults to lay their eggs. Such effect did not happen in ponds where fish was present likely because fish increased predation pressure upon dragonflies as other suitable prey became scarce. This might also explain why *Orthemis* dragonflies consistently had higher abundances in ponds with fish in low isolation treatments, compared to fishless ponds. It is possible that the higher availability of more suitable prey (*i.e.* dytiscid beetles and notonectids) in low isolation decreased predation rate upon *Orthemis*, allowing it to have a greater abundance in ponds with fish. Indeed, some dragonfly species are known to exhibit different vulnerability to predation depending on body size and antipredatory behavior, allowing them to coexist with fish (Johnson 1991, McPeek 1998, De Marco et al. 1999, Johansson 2000, Hopper 2001, McCauley 2008).

The reduction in total abundance of predatory insects across the isolation gradient indeed had effects that are analogous to a trophic cascade, causing the abundance of herbivores and detritivores to increase, but this effect was only clear in fishless ponds both considering total abundance and community structure patterns. We argue that the most plausible explanation for this pattern lies in the fact that, because the general abundance of predatory insects decreases in more isolated ponds, predation pressure by fish upon herbivores and detritivores may also increase with isolation, again, counteracting the indirect positive effect of the reduction in the abundance of predatory insects.

Our main hypothesis was that the effect of fish would be stronger as spatial isolation increases, causing fish and fishless communities to have greater differences in structure at more isolated ponds (Fig. 1c). We did find this pattern (Fig. 3), however, we initially predicted it would be a consequence of a reinforced effect of fish by spatial isolation. As discussed above, herbivores and detritivores did not have similar responses to isolation and presence of generalist fish. Instead, when generalist fish was present, it dampened the indirect positive effects of isolation. Additionally, not all predatory insects were limited by dispersal, which made the ones that are good dispersers to thrive in more isolated ponds when fish was absent. Therefore, we argue that the greater differences between communities with and without fish in more isolated ponds are because the presence of fish changes the net effects of spatial isolation on community structure. When fish is absent, spatial isolation generally favors the abundance of aquatic insects with higher dispersal rates, regardless of their trophic level. When fish is present, spatial isolation has negative effects on the abundance of predatory insects, and a weaker positive effect on the abundance of herbivores and detritivores. These different paths of change in structure were the cause of greater differences among fish and fishless ponds in our higher isolation treatments. Furthermore, those patterns could be generated by either consumptive (*i.e.* direct predation upon available prey) or non-consumptive effects of predators upon preys, like habitat selection (*i.e.* avoidance of ponds with fish or high density of predatory insects; see Binckley and Resetarits 2005, Blaustein et al. 2005, Resetarits 2005), still, it does not change the general conclusions of this work, as both processes represent a negative local selective pressure of predators upon preys.

Finally, our results diverge both from simplistic metacommunity models, which consider similar dispersal rates and single trophic levels (Mouquet and Loreau 2003, Leibold et al. 2004, Leibold and Chase 2018), and slightly more complex ones considering variation in dispersal rates (Vellend et al. 2014). We argue that the complexity of our study system, both in terms of interspecific variability in dispersal rates and complex indirect effects involved in trophic interactions, are the main causes of such divergence. Nonetheless, we were able to show that spatial isolation can cause effects that are similar to trophic cascades on freshwater communities, and that the presence of a generalist top predator can dampen those effects. Our results also highlight the importance of considering interspecific variation in dispersal rates and multiple trophic levels to better understand the processes generating community and metacommunity structures in spatially structured landscapes (see Van De Meutter et al. 2007, Vellend et al. 2014, Hill et al. 2017, Guzman et al. 2019).

## Acknowledgments

We thank the EESB staff for assistance in pond construction and Luis Vicente Cavalaro, Bianca Valente, Fernanda Simioni, Débora Negrão, Jessika Akane and Suzana Marte for assistance in the community sampling surveys. We thank Tadeu Siqueira and Paulo Inácio Prado for conceptual and statistical advice, Victor Saito and Erika Shimabukuro for assistance in the identification of aquatic insects, Renata Pardini and Daniel Lahr for providing lab and office space, Paulo Roberto Guimarães Junior and Paulo De Marco Júnior for insightful comments on early versions of the manuscript; The members of the Schiesari lab and Leibold lab for criticism; and Giselda Durigan for introducing us to the EESB. This study was funded by The São Paulo Research Foundation (FAPESP, grant #2014/16320-7, Carlos Navas PI, Luis Schiesari Associate Researcher; FAPESP, grant #2015/18790-3; Luiz Antonio Martinelli PI, Luis Schiesari Co-PI) and National Council for Scientific and Technological Development (CNPq, grant #458796/2014-0). RMP was supported by Ph.D. fellowships from FAPESP (grants #2017/04122-4 and #2018/07714-2) and Coordination of Superior Level Staff Improvement (CAPES). Experiments were conducted in EESB under the authorization from Instituto Florestal (COTEC 553/2017) and collection permits from Instituto Chico Mendes de Conservação da Biodiversidade (ICMBIo 17559-6), following protocols approved by the Research Ethics Committee of the School of Arts, Sciences and Humanities of the University of São Paulo (CEUA 003/2016).

