## Supplemental Information for "Top predator introduction changes the effects of spatial isolation on freshwater community structure"

### Index

**Appendix S1**

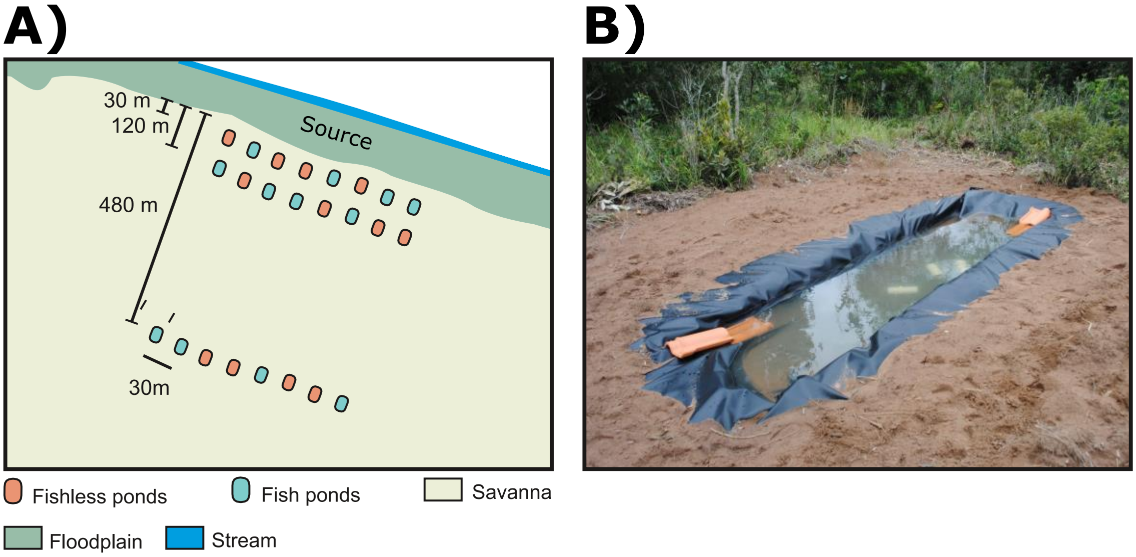

**Figure S1.1.** A. Diagram of the experimental layout. B. Representative photo of one of our experimental units.

#### Appendix S2

**Table S2.1.** Macroinvertebrate dispersal abilities are poorly known, especially in the Neotropical region. To support our decisions in experimental design we conducted a brief, non-exhaustive review of the distances at which macroinvertebrate families that colonized our mesocosms experience negative effects of isolation. It is important to mention that information about dispersion ability are scarce, imprecise, and many times contradictory since they mostly rely on abundance data, which are likely to be affected by many other factors. Still, many of the negative effects of spatial isolation on aquatic macroinvertebrates can be found within the range of distances manipulated in our experiment (30-120-480 m).

| | **Distance of minimum isolation effect (m)** | **Reference** |
|---|---:|---:|
| Dytiscidae | 50 | Shulman and Chase 2007 |
| Hydrophilidae | ? | ? |
| Dryopidae | ? | ? |
| Ceratopogonidae | ? | ? |
| Chaoboridae | 50 | van de Meutter et al. 2006; Chase and Shulman 2009 |
| Chironomidae | >500 | Delettre and Morvan 2000; van de Meutter et al. 2006; Shulman and Chase 2007; Muehlbauer et al. 2014 |
| Chironomidae (Tanypodinae) | ~ 100 | Delettre and Morvan 2000; van de Meutter et al. 2006 |
| Culicidae | >1500 | Chase and Shulman 2009 |
| Baetidae | ? | Delettre and Morvan 2000; Muehlbauer et al. 2014 |
| Caenidae | ~ 160[1] | Muehlbauer et al. 2014 |
| Polimitarcyidae | ~ 160[1] | Muehlbauer et al. 2014 |
| Corixidae | ? | - |
| Notonectidae. | ~ 300 | Wilcox 2001; Trekels et al. 2011 |
| Veliidae | ? | - |
| Gerridae | ? | - |
| Libellulidae | >1000 | McCauley 2006 |
| Coenagrionidae | ~ 300 | Purse et al. 2003; van de Meutter et al. 2006 |

**Table S2.2.** Macroinvertebrate traits that contribute to dispersal rates. Voltinism refers to the number of generations within a year. Families were classified as being Semivoltine (< 1 generation/year), Univoltine (1 generation/year) or Multivoltine (> 1 generation/year). Development was classified as either Slow or Fast larval development, and Seasonal or Nonseasonal. Adult life span was classified as Very Short (< 1 week), Short (< 1 month) or Long (> 1 month). Adult ability to exit pond refers to the possibility of adults to disperse among aquatic habitats other than in the moment of emergence. Adult flying strength were classified as Weak (cannot fly into light breeze) or Strong (can fly into light breeze). Taxa that are multivoltine and that have fast development, short adult life span, ability to exit ponds, and higher-flying strength are thought to have higher dispersal rates. Note, however, that this general classification should be regarded with caution as it is mostly based on temperate macroinvertebrates.

| | Voltinism | Development | Adult Life Span | Adult ability to exit pond | Adult flying strength | Reference |
|---|---|---|---|---|---|---|
| Dytiscidae | Semivoltine | Slow seasonal | Long | YES | Strong | Poff et al. 2006 |
| Hydrophilidae | Semivoltine | ? | ? | NO | ? | Hosseinie 1995; Resetarits 2001 |
| Dryopidae | Semivoltine | Slow seasonal and Nonseasonal | Long | NO | Weak | Poff et al. 2006 |
| Ceratopogonidae | Univoltine | Fast Seasonal | Very short | NO | Weak | Poff et al. 2006 |
| Chaoboridae | Uni or Multivoltine | ? | Very short | NO | Weak | Thorp & Covich 2009 |
| Chironomidae | Multivoltive | Fast Seasonal | Very short | NO | Weak | Poff et al. 2006; Hamada et al. 2014 |
| Chironomidae (Tanypodinae) | Multivoltive | Fast Seasonal | Very short | NO | Weak | Poff et al. 2006; Hamada et al. 2014 |
| Culicidae | Multivoltive | Fast Seasonal | Very short | NO | Weak | Ciota et al. 2014 |
| Baetidae | Multivoltive | Fast Seasonal | Very short | NO | Weak | Poff et al. 2006 |

| | | | | | | |
|---|---|---|---|---|---|---|
| Caenidae | Multivoltive | Slow seasonal | Very short | NO | Weak | Poff et al. 2006 |
| Polimitarcyidae | Univoltine | Slow seasonal | Very short | NO | Weak | Poff et al. 2006 |
| Corixidae | Multivoltive | Fast Seasonal | Long | YES | Strong | Poff et al. 2006 |
| Notonectidae. | Uni or Multivoltine | ? | ? | YES | ? | Thorp and Covich 2009 |
| Veliidae | Multivoltive | Fast Seasonal | Long | YES | Weak | Poff et al. 2006 |
| Gerridae | Univoltine | Fast Seasonal | Long | YES | Weak | Poff et al. 2006 |
| Libellulidae | Semivoltine | Slow seasonal | Long | NO | Strong | Poff et al. 2006 |
| Coenagrionidae | Univoltine | Slow seasonal | Short | NO | Weak | Poff et al. 2006 |

**Appendix S3**

**Table S3.1.** Dates of fish additions to each pond and the most conservative possible estimation of the number of days that each pond remained fishless (including fish and fishless treatments for comparison).

| Pond ID | Fish Treatment | Isolation Treatment | Fish addition | Confirmation of absence of fish | Maximum number of days that pond remained fishless | | |
|---|---|---|---|---|---|---|---|
| | | | | | Before the First Survey | Between First and Second Survey | Between Second and Third Survey |
| **A1** | present | 30 m | 30-Jan / 24-Feb / 27-Mar | 24-Feb / 26-Mar | 16 | 31 | 0 |
| **A2** | present | 30 m | 30-Jan / 01-Feb | 31-Jan | 1 | 0 | 0 |
| **A3** | absent | 30 m | | | 24 | 31 | 28 |
| **A4*** | present | 30 m | 30-Jan | | 16 | 0 | 0 |
| **A5** | absent | 30 m | | | 24 | 31 | 28 |
| **A6** | absent | 30 m | | | 24 | 31 | 28 |
| **A7** | present | 30 m | 30-Jan / 01-Feb / 27-Mar | 31-Jan / 01-Fev | 0 | 31 | 0 |
| **A8** | absent | 30 m | | | 24 | 31 | 28 |
| **B1** | absent | 120 m | | | 24 | 31 | 28 |
| **B2** | absent | 120 m | | | 24 | 31 | 28 |
| **B3*** | present | 120 m | 30-Jan | | 0 | 0 | 0 |
| **B4** | present | 120 m | 30-Jan | | 0 | 0 | 0 |
| **B5** | absent | 120 m | | | 24 | 31 | 28 |
| **B6** | present | 120 m | 30-Jan | | 0 | 0 | 0 |
| **B7** | absent | 120 m | | | 24 | 31 | 28 |
| **B8** | present | 120 m | 30-Jan / 01-Feb | 31-Jan | 1 | 0 | 0 |
| **C1** | present | 480 m | 30-Jan | | 0 | 0 | 0 |
| **C2** | absent | 480 m | | | 24 | 31 | 28 |
| **C3*** | absent | 480 m | | | 24 | 31 | 28 |
| **C4*** | present | 480 m | 30-Jan /07-Feb | 06-Feb | 6 | 0 | 0 |
| **C5** | absent | 480 m | | | 24 | 31 | 28 |

| | | | | | | | |
|---|---|---|---|---|---|---|---|
| **C6** | absent | 480 m | | | 24 | 31 | 28 |
| **C7** | present | 480 m | 30-Jan | | 0 | 0 | 0 |
| **C8** | present | 480 m | 30-Jan / 01-Feb | 31-Jan | 1 | 0 | 0 |

*Samples excluded in the third survey

#### Appendix S4

**Table S4.1.** Pilot laboratory experiment in which we offered four individuals of a variety of vertebrate and invertebrate prey common at our study site to each of eight Redbreast Tilapias in individual aquaria.

| Date | Average room temperature | Fish ID | Taxa | Number of individuals left after: | | | | | |
|---|---|---|---|---|---|---|---|---|---|
| | | | | 0 (min) | 30 (min) | 60 (min) | 120 (min) | 240 (min) | 1440 (min) |
| **09-Jan/2017** | 26.7 | 1 | *Phalloceros* sp. | 4 | 4 | 4 | 4 | 4 | 4 |
| **10-Jan/2017** | 27.4 | 1 | Zigoptera | 4 | 2 | 0 | 0 | 0 | 0 |
| **11-Jan/2017** | 29.2 | 1 | Nepidae | 4 | 3 | 2 and 1* | 2 and 1* | 2 and 1* | 2 and 1* |
| **09-Jan/2017** | 26.7 | 2 | *Scinax* sp. | 4 | 0 | 0 | 0 | 0 | 0 |
| **10-Jan/2017** | 27.4 | 2 | Anisoptera | 4 | 0 | 0 | 0 | 0 | 0 |
| **11-Jan/2017** | 29.2 | 2 | *Phalloceros* sp. | 4 | 4 | 4 | 3 | 3 | 0 |
| **12-Jan/2017** | 29.2 | 2 | Nepidae (Large) | 4 | 4 | 4 | 4 | 4 | 4 |
| **13-Jan/2017** | - | 2 | *Aedes* | 4 | 0 | 0 | 0 | 0 | 0 |
| **09-Jan/2017** | 26.7 | 3 | Anisoptera | 4 | 4 | 4 | 0 | 0 | 0 |
| **10-Jan/2017** | 27.4 | 3 | Nepidae (Large) | 4 | 3 | 2 and 1* | 2 and 1* | 2 and 1* | 1 |
| **11-Jan/2017** | 29.2 | 3 | *Phalloceros* sp. | 4 | 4 | 4 | 4 | 4 | 0 |
| **12-Jan/2017** | 29.2 | 3 | *Scinax* sp. | 4 | 0 | 0 | 0 | 0 | 0 |
| **13-Jan/2017** | - | 3 | Belostomatidae | 4 | 0 | 0 | 0 | 0 | 0 |
| **09-Jan/2017** | 26.7 | 4 | *Scinax* sp. | 4 | 0 | 0 | 0 | 0 | 0 |
| **10-Jan/2017** | 27.4 | 4 | *Phalloceros* sp. | 4 | 0 | 0 | 0 | 0 | 0 |
| **11-Jan/2017** | 29.2 | 4 | Anisoptera | 4 | 0 | 0 | 0 | 0 | 0 |
| **12-Jan/2017** | 29.2 | 4 | Zigoptera | 4 | 0 | 0 | 0 | 0 | 0 |
| **09-Jan/2017** | 26.7 | 5 | Nepidae (Small) | 4 | 0 | 0 | 0 | 0 | 0 |
| **10-Jan/2017** | 27.4 | 5 | Beetle | 4 | 4 | 4 | 4 | 4 | 4 |
| **11-Jan/2017** | 29.2 | 5 | *Scinax* sp. | 4 | 0 | 0 | 0 | 0 | 0 |
| **12-Jan/2017** | 29.2 | 5 | Anisoptera | 4 | 0 | 0 | 0 | 0 | 0 |

| | | | | | | | | | |
|---|---|---|---|---|---|---|---|---|---|
| **13-Jan/2017** | - | 5 | *Phalloceros* sp. | 4 | 4 | 4 | 4 | 0 | 0 |
| **09-Jan/2017** | 26.7 | 6 | Zigoptera | 4 | 0 | 0 | 0 | 0 | 0 |
| **10-Jan/2017** | 27.4 | 6 | *Phalloceros* sp. | 4 | 4 | 4 | 4 | 4 | 0 |
| **11-Jan/2017** | 29.2 | 6 | Anisoptera | 4 | 0 | 0 | 0 | 0 | 0 |
| **09-Jan/2017** | 26.7 | 7 | *Phalloceros* sp. | 4 | 4 | 4 | 4 | 4 | 0 |
| **10-Jan/2017** | 27.4 | 7 | Nepidae (Large) | 4 | 4 | 4 | 4 | 4 | 3* |
| **11-Jan/2017** | 29.2 | 7 | Anisoptera | 4 | 0 | 0 | 0 | 0 | 0 |
| **09-Jan/2017** | 26.7 | 8 | Anisoptera | 4 | 0 | 0 | 0 | 0 | 0 |
| **10-Jan/2017** | 27.4 | 8 | Nepidae (Small) | 4 | 0 | 0 | 0 | 0 | 0 |
| **11-Jan/2017** | 29.2 | 8 | Zigoptera | 4 | 0 | 0 | 0 | 0 | 0 |
| **12-Jan/2017** | 29.2 | 8 | *Phalloceros* sp. | 4 | 3 | 3 | 2 | 0 | 0 |

*Numbers with an asterisk symbol represent half of an individual, meaning that the other half was eaten by the fish.

**Appendix S5**

Due to inevitable practical constraints, one limitation of our experimental design is that experimental units were not fully spatially isolated from each other. That is, one might hypothesize that as local insect populations build up over time and emerge from ponds, they may affect the local dynamics of neighboring ponds. We, therefore, tested whether the spatial configuration of the experimental ponds could explain the community structures we observed at the end of our experiment (*i.e.* last survey). To do so, we computed artificial spatial variables using distance-based Moran's Eigenvector Maps (MEMs). MEMs are computed from a truncated Euclidian distance matrix built using geographical coordinates of the experimental units (Borcard and Legendre 2002, Dray et al. 2006, 2012). In this case, values above the truncation distance are multiplied by 4 to emphasize the relationship between experimental units that are closer together. Normally the truncation distance is set to the largest necessary distance for all sample units to stay connected (*i.e.* minimum spanning tree). However, because here we wanted to emphasize fine spatial patterns we set our truncation distance to 60m, which allows for any pond to have a disproportionally larger effect on ponds located at a maximum of 60m distance (i.e., any pond could be influenced by 4 neighboring ponds; Figure 1).

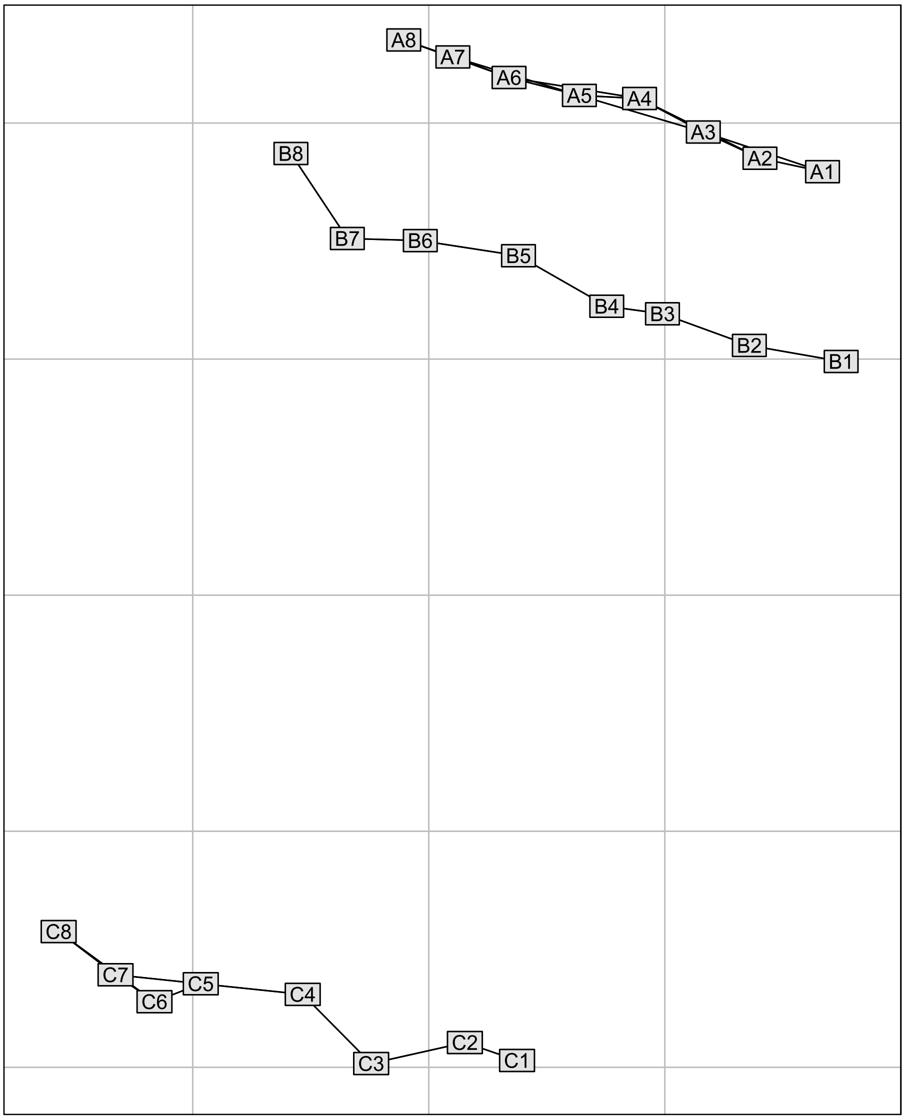

**Figure S5.1.** The spatial configuration of our experimental units. Each rectangle represents one mesocosm plotted according to their geographic coordinates. Only ponds that are within a 60 m distance from each other are connected, meaning that they are more likely to affect each other through spatial dynamics.

We then performed a Principal Coordinate Analysis (PCoA) on the truncated distance matrix and select the axes with positive eigenvalues to represent the artificial spatial variables. Axes with negative eigenvalues represent negative spatial autocorrelation (Dray et al. 2006), which would mean that ponds closer together are less similar to each other them ponds that are further apart. The first axes from the PCoA represent broader spatial patterns, while the last ones represent finer ones, which is what we are mostly interested in (Figure 2).

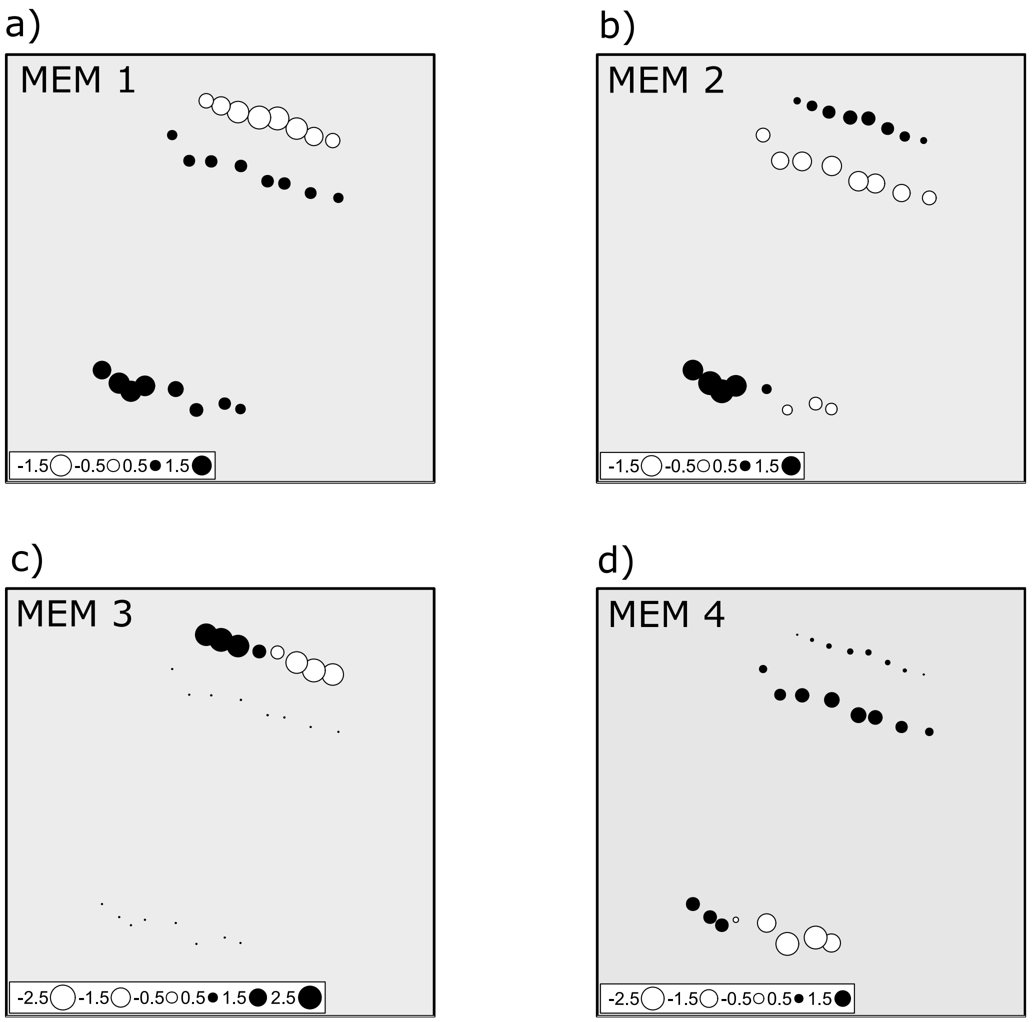

**Figure S5.2.** Plots of the artificial spatial variables (MEMs) generated by the PCoA on the truncated distance matrix. Each circle represents one experimental unit positioned according to its geographic coordinate. The size and color of the circles are proportional to the value attributed in each MEM. In this case, circles of similar colors and sizes are similar to each other.

With the artificial spatial variables in hand, we first performed sequential likelihood ratio tests to test if any of the four spatial variables recovered from the PCoA provided a significantly better fit to our community data (Table 1). As expected, we found that only the addition of the first axis provided a better fit to our data, which mostly describes the isolation distances that we manipulated (Figure S1.2a). Then we repeated this likelihood ratio tests but now after accounting to variation in community structure explained by our fish and isolation treatments (Table 2). We found that none of the MEMs significantly explained variation in community structure. This means that our manipulated treatments already accounted for any spatial effects described by any of the MEMs. Therefore, we conclude that any spatial processes caused by the proximity of experimental ponds at any given isolation distance had a minor, if any, importance in the structure of the experimental communities.

**Table S5.1.** Sequential likelihood ratio tests for the effect of each of the MEMs on community patterns.

| Model | Residual Df. | Df. diff | Dev. | p |
|---|---|---|---|---|
| **MEM1** | **18** | **1** | **40.33** | **0.037** |
| MEM2 | 17 | 1 | 19.87 | 0.536 |
| MEM3 | 16 | 1 | 25.72 | 0.345 |
| MEM4 | 15 | 1 | 37.38 | 0.221 |

**Table S5.2.** Sequential likelihood ratio tests including the MEMs after the treatment factors.

| Model | Residual Df. | Df. diff | Dev. | p |
|---|---|---|---|---|
| **Fish** | **18** | **1** | **49.09** | **0.018** |
| Isolation | 16 | 2 | 72.96 | 0.054 |
| **Fish : Isolation** | **14** | **2** | **91.12** | **0.015** |
| MEM1 | 13 | 1 | 39.43 | 0.143 |
| MEM2 | 12 | 1 | 22.54 | 0.717 |
| MEM3 | 11 | 1 | 31.73 | 0.352 |
| MEM4 | 10 | 1 | 62.40 | 0.100 |

#### Appendix S6

**Table S6.1.** Abundance and traits (trophic level, maximum recorded body volume in experiment) of taxa colonizing experimental ponds.

| Order | Family | Subfamily | Taxa | Abbreviation | Volume of largest individual (mm³) | Wet mass of largest individual (mg) | Total Abundance | Trophic Level | Reference to Trophic Level |
|---|---|---|---|---|---|---|---|---|---|
| **Coleoptera** | Dytiscidae | | *Rhantus* | Rha | 387.52 | 36.28 | 14 | Predator | Ramírez and Gutiérrez-Fonseca 2014 |
| **Coleoptera** | Dytiscidae | | *Thermonectus* | The | 0.51 | 0.06 | 3 | Predator | Ramírez and Gutiérrez-Fonseca 2014 |
| **Coleoptera** | Dytiscidae | | *Derovatellus* | Der | 2.60 | 0.2 | 2 | Predator | Ramírez and Gutiérrez-Fonseca 2014 |
| **Coleoptera** | Dytiscidae | | *Hydaticus* | Hda | 210.15 | 27 | 1 | Predator | Ramírez and Gutiérrez-Fonseca 2014 |
| **Coleoptera** | Hydrophilidae | | *Berosus* | Ber | 5.02 | 19.66 | 4 | Predator | Ramírez and Gutiérrez-Fonseca 2014 |
| **Coleoptera** | Hydrophilidae | | *Tropisternus* | Tro | 231.38 | 1.02 | 2 | Predator | Ramírez and Gutiérrez-Fonseca 2014 |
| **Coleoptera** | Noteridae | | *Hydrocanthus* | Hdr | 3.14 | 0.42 | 1 | Predator | Ramírez and Gutiérrez-Fonseca 2014 |
| **Diptera** | Ceratopogonidae | | Ceratopogonidae | Cer | 0.65 | 0.24 | 6 | Herbivore/ Detritivore** | Aussel and Linley 1994, Ramírez and Gutiérrez-Fonseca 2014 |
| **Diptera** | Chaoboridae | | *Chaoborus* | Cha | 4.29 | 0.31 | 68 | Herbivore/ Detritivore ** | Arcifa 2000, Ramírez and Gutiérrez-Fonseca 2014 |
| **Diptera** | Chironomidae | Chironominae | Tanytarsini | Tan | 4.44* | 0.72* | 7214 | Herbivore/ Detritivore | Ramírez and Gutiérrez-Fonseca 2014 |
| **Diptera** | Chironomidae | Chironominae | *Goeldchironomus* | Goe | 4.44* | 0.72* | 1455 | Herbivore/ Detritivore | Ramírez and Gutiérrez-Fonseca 2014 |
| **Diptera** | Chironomidae | Chironominae | *Polypedilum* | Pol | 4.44* | 0.72* | 1436 | Herbivore /Detritivore | Ramírez and Gutiérrez-Fonseca 2014 |
| **Diptera** | Chironomidae | Chironominae | *Chironomus* | Chi | 4.44* | 0.72* | 534 | Herbivore/ Detritivore | Ramírez and Gutiérrez-Fonseca 2014 |
| **Diptera** | Chironomidae | Chironominae | *Asheum* | Ash | 4.44* | 0.72* | 399 | Herbivore/ Detritivore | Ramírez and Gutiérrez-Fonseca 2014 |
| **Diptera** | Chironomidae | Chironominae | *Caladomyia* | Cla | 4.44* | 0.72* | 38 | Herbivore/ Detritivore | Ramírez and Gutiérrez-Fonseca 2014 |

| | | | | | | | | | |
|---|---|---|---|---|---|---|---|---|---|
| **Diptera** | Chironomidae | Chironominae | *Apedilum* | Ape | 4.44* | 0.72* | 4 | Herbivore/ Detritivore | Ramírez and Gutiérrez-Fonseca 2014 |
| **Diptera** | Chironomidae | Chironominae | *Beardius* | Bea | 4.44* | 0.72* | 3 | Herbivore/ Detritivore | Ramírez and Gutiérrez-Fonseca 2014 |
| **Diptera** | Chironomidae | Chironominae | *Parachironomus* | Par | 4.44* | 0.72* | 3 | Herbivore/ Detritivore | Ramírez and Gutiérrez-Fonseca 2014 |
| **Diptera** | Chironomidae | Tanytarsinae | *Ablabesmyia* | Abl | 3.28* | 0.33* | 363 | Herbivore/ Detritivore ** | Henriques-Oliveira et al. 2003, Ramírez and Gutiérrez-Fonseca 2014 |
| **Diptera** | Chironomidae | Tanytarsinae | *Larsia* | Lar | 3.28* | 0.33* | 100 | Herbivore/ Detritivore ** | Henriques-Oliveira et al. 2003, Ramírez and Gutiérrez-Fonseca 2014 |
| **Diptera** | Chironomidae | Tanytarsinae | *Labrundinia* | Lab | 3.28* | 0.33* | 9 | Herbivore/ Detritivore ** | Henriques-Oliveira et al. 2003, Ramírez and Gutiérrez-Fonseca 2014 |
| **Diptera** | Culicidae | | *Culex* | Cul | 2.42 | 0.42 | 707 | Herbivore/ Detritivore | Ramírez and Gutiérrez-Fonseca 2014 |
| **Ephemeroptera** | Baetidae | | *Callibaetis* | Cal | 12.56 | 0.95 | 14 | Herbivore/ Detritivore | Ramírez and Gutiérrez-Fonseca 2014 |
| **Ephemeroptera** | Caenidae | | *Caenis* | Cae | 17.53 | 0.26 | 155 | Herbivore/ Detritivore | Ramírez and Gutiérrez-Fonseca 2014 |
| **Ephemeroptera** | Polymitarcyidae | | *Campsurus* | Cam | 68.81 | 5.01 | 10 | Herbivore/Detritivore | Ramírez and Gutiérrez-Fonseca 2014 |
| **Hemiptera** | Corixidae | | *Tenagobia* | Ten | 1.72 | 0.09 | 3 | Herbivore/Detritivore | Ramírez and Gutiérrez-Fonseca 2014 |
| **Hemiptera** | Corixidae | | *Heterocorixa* | Het | 16.73 | 1.82 | 1 | Herbivore/Detritivore | Ramírez and Gutiérrez-Fonseca 2014 |
| **Hemiptera** | Gerridae | | *Rheumatobates* | Rhe | 38.00 | 4.37 | 1 | Predator | Ramírez and Gutiérrez-Fonseca 2014 |
| **Hemiptera** | Naucoridae | | *Ctenipocoris* | Cte | 27.42 | 2.04 | 1 | Predator | Ramírez and Gutiérrez-Fonseca 2014 |
| **Hemiptera** | Notonectidae | | *Buenoa* | Bue | 34.43 | 4.77 | 139 | Predator | Ramírez and Gutiérrez-Fonseca 2014 |
| **Hemiptera** | Notonectidae | | *Notonecta* | Not | 118.11 | 13.05 | 6 | Predator | Ramírez and Gutiérrez-Fonseca 2014 |
| **Hemiptera** | Veliidae | | *Microvelia* | Mic | 3.44 | 0.3 | 1097 | Predator | Ramírez and Gutiérrez-Fonseca 2014 |
| **Odonata** | Coenagrionidae | | *Oxyagrion* | Oxy | 8.74 | 0.68 | 5 | Predator | Ramírez and Gutiérrez-Fonseca 2014 |
| **Odonata** | Libellulidae | | *Erythrodiplax* | Ery | 163.48 | 4.96 | 465 | Predator | Ramírez and Gutiérrez-Fonseca 2014 |

| | | | | | | | | |
|---|---|---|---|---|---|---|---|---|
| **Odonata** | Libellulidae | *Pantala* | Pan | 1054.38 | 29 | 349 | Predator | Ramírez and Gutiérrez-Fonseca 2014 |
| **Odonata** | Libellulidae | *Orthemis* | Ort | 52.50 | 2.6 | 87 | Predator | Ramírez and Gutiérrez-Fonseca 2014 |

*We used the same volume and wet mass value for all Chironominae and Tanytarsinae taxa.

**Insects that also have predatory behavior recorded but mostly consume plankton (either phytoplankton or zooplankton) or only small amounts or parts of the other organisms considered in our study.

**Appendix S7**

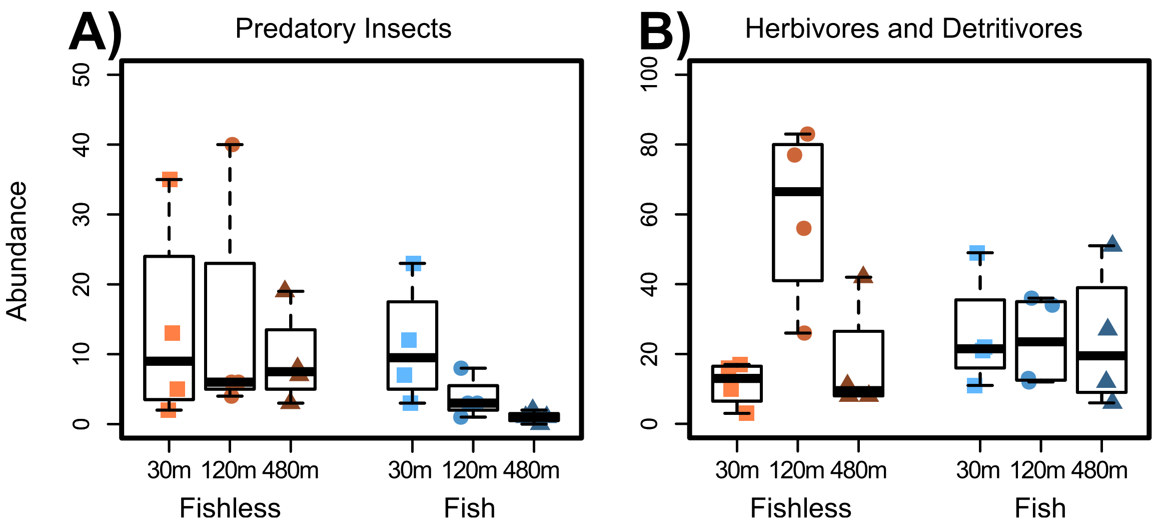

**Figure S7.1.** Total abundance of predatory insects (A) and herbivores and detritivores (B) in the first survey.

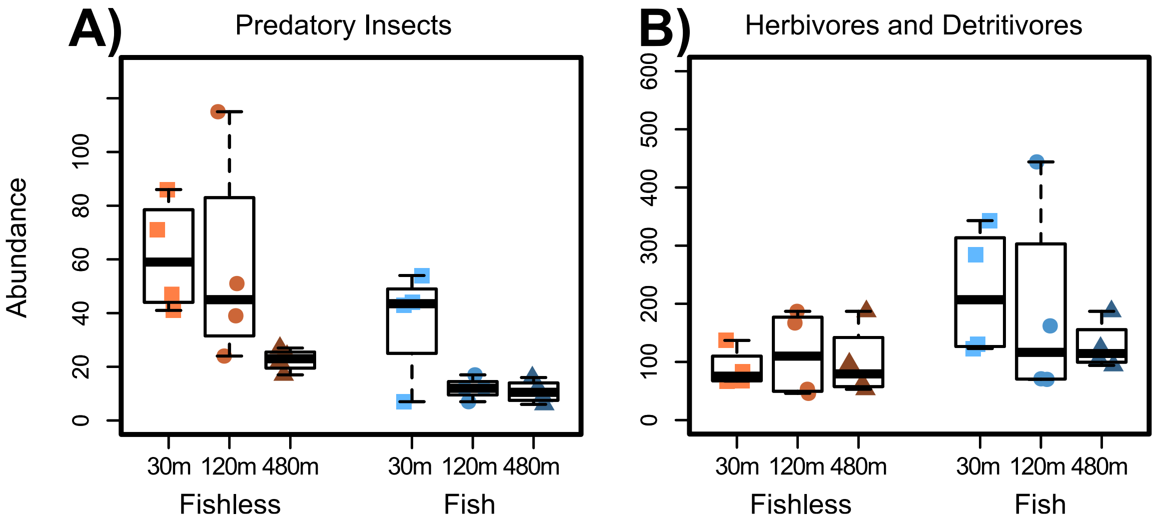

**Figure S7.2.** Total abundance of predatory insects (A) and herbivores and detritivores (B) in the second survey.

**Appendix S8**

**Figure S8.1.** Model-based unconstrained ordinations showing pond communities (symbols) and species (bubbles) in each of the three sampling surveys. Yellow bubbles are predatory-insects and green bubbles are herbivores and detritivores. Size of bubbles are proportional to body size of each taxa (the volume of the largest individual of each species in a log-scale). A – First sampling survey; B – Second sampling survey; Abbreviations of names of taxa provided in Appendix S6.

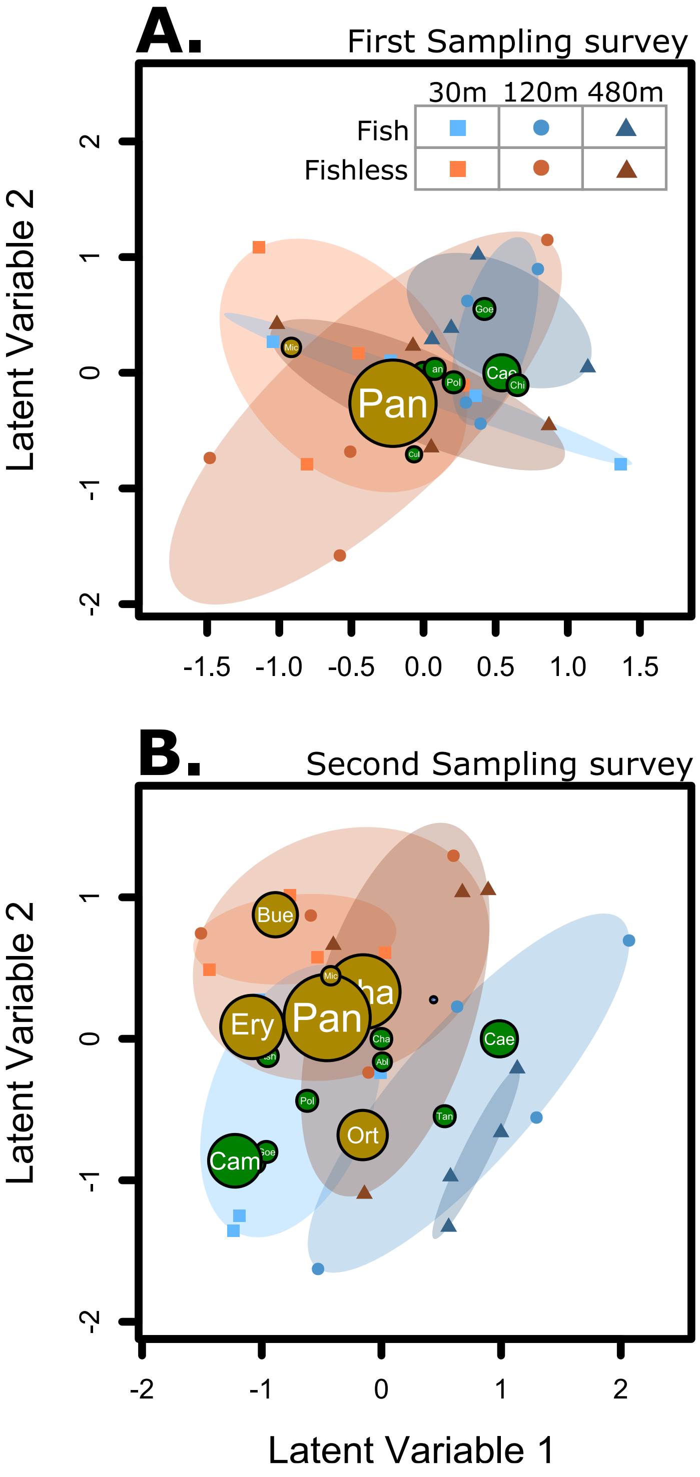

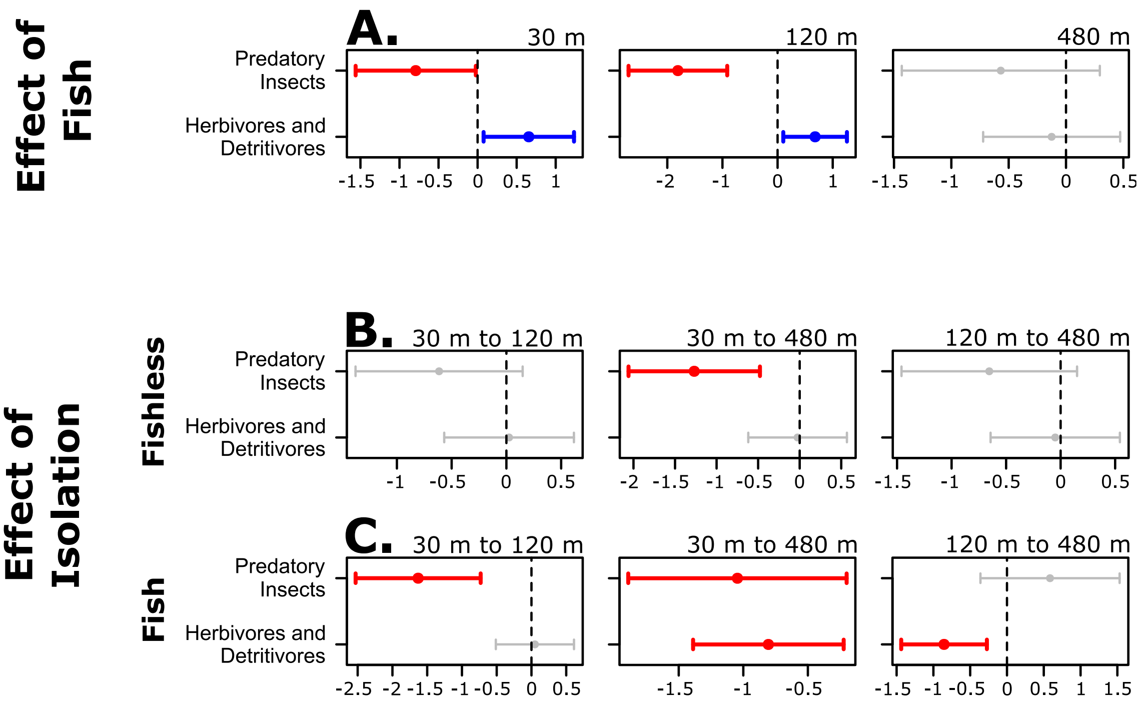

**Figure S8.2.** 95% Confidence intervals for the effect of fish and isolation on abundance of predators and herbivores and detritivores when comparing pairs of treatments for the second survey of the experiment. Confidence intervals not crossing the zero hatched line were considered significant effects and colored; blue bars represent an increase and red bars a decrease in abundance from the reference treatment. A are effects of the presence of fish in each isolation treatment. B are effects of isolation in fishless ponds and C in ponds with fish. In each of the C and B we show effects of increasing isolation from 30 to 120 m, from 30 to 480 m, and from 120 m to 480 m.

**Table S8.1.** Increase (positive values) or decrease (negative values) in the effect of fish from one level of isolation to another measured by the difference in distance between the centroids of each treatment in a model-based unconstrained ordination. Bold lines represent significative increase or decrease in distance values. P values were adjusted for false discovery ratio.

| | **difference in effect of fish** | **p value** | **adj. p value** |
|---|---|---|---|
| 1st Sampling Survey | | | |
| 30 m to 120 m | 0.410588 | 0.5301 | 0.8565 |
| 30 m to 480 m | 0.040839 | 0.949 | 0.949 |
| 120 m to 480 m | -0.369749 | 0.571 | 0.8565 |
| 2nd Sampling Survey | | | |
| 30 m to 120 m | 0.27635 | 0.6525 | 0.9798 |
| 30 m to 480 m | 0.003335 | 0.996 | 0.996 |
| 120 m to 480 m | -0.273015 | 0.6532 | 0.9798 |
| 3rd Sampling Survey | | | |
| 30 m to 120 m | 1.251939 | 0.0654 | 0.0981 |
| **30 m to 480 m** | **1.628767** | **0.0126** | **0.0378** |
| 120 m to 480 m | 0.376828 | 0.6011 | 0.6011 |

#### Appendix S9

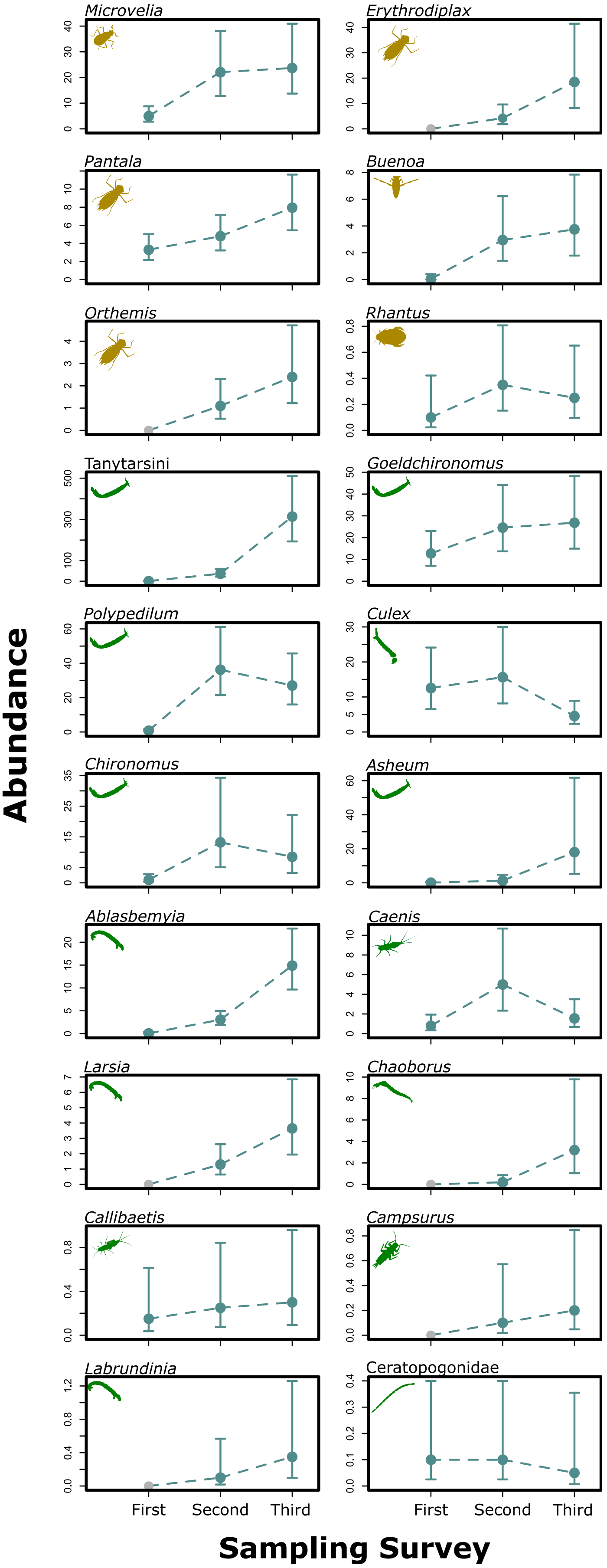

**Figure S9.1.** Effect of time on species abundances according to maximum likelihood estimates of abundance and their 95% confidence interval for Model 1 in Table 1. Grey symbols indicate absolute absence (zero abundance) of a taxon in a treatment. More information about the estimated effects are provided in Appendix 9.

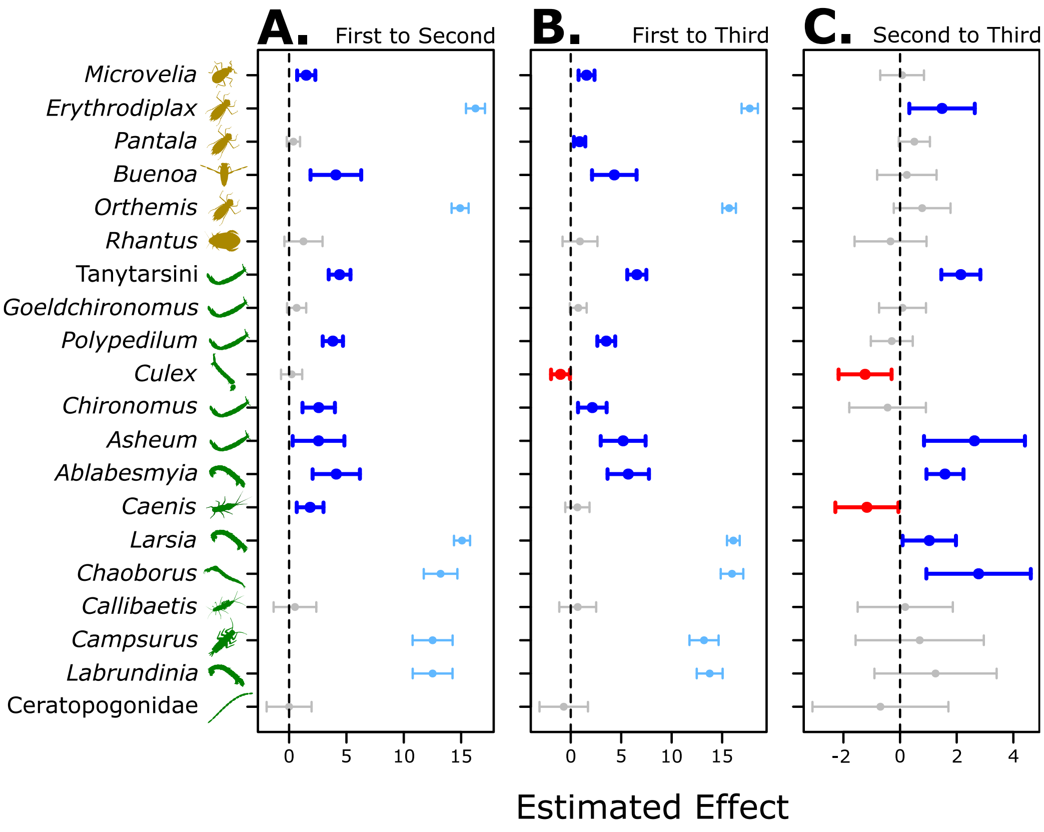

**Figure S9.2.** Confidence intervals for the effect of time on the abundance of each taxon, predatory insects (yellow), and herbivores and detritivores (green). Taxa are ordered from most (top) to less abundant (bottom) according to trophic level (predators and herbivores and detritivores, respectively). Bars with 95% confidence intervals not crossing the zero-line are considered significant and colored. Blue bars represent an increase in abundance and red bars a decrease in abundance relative to the reference treatment. Light blue bars represent taxa that were absent from the reference treatment. A – Effect of moving from the first to the second sampling survey; B – Effect of moving from the first to the third sampling survey; C – Effect of moving from the second to the third sampling survey.

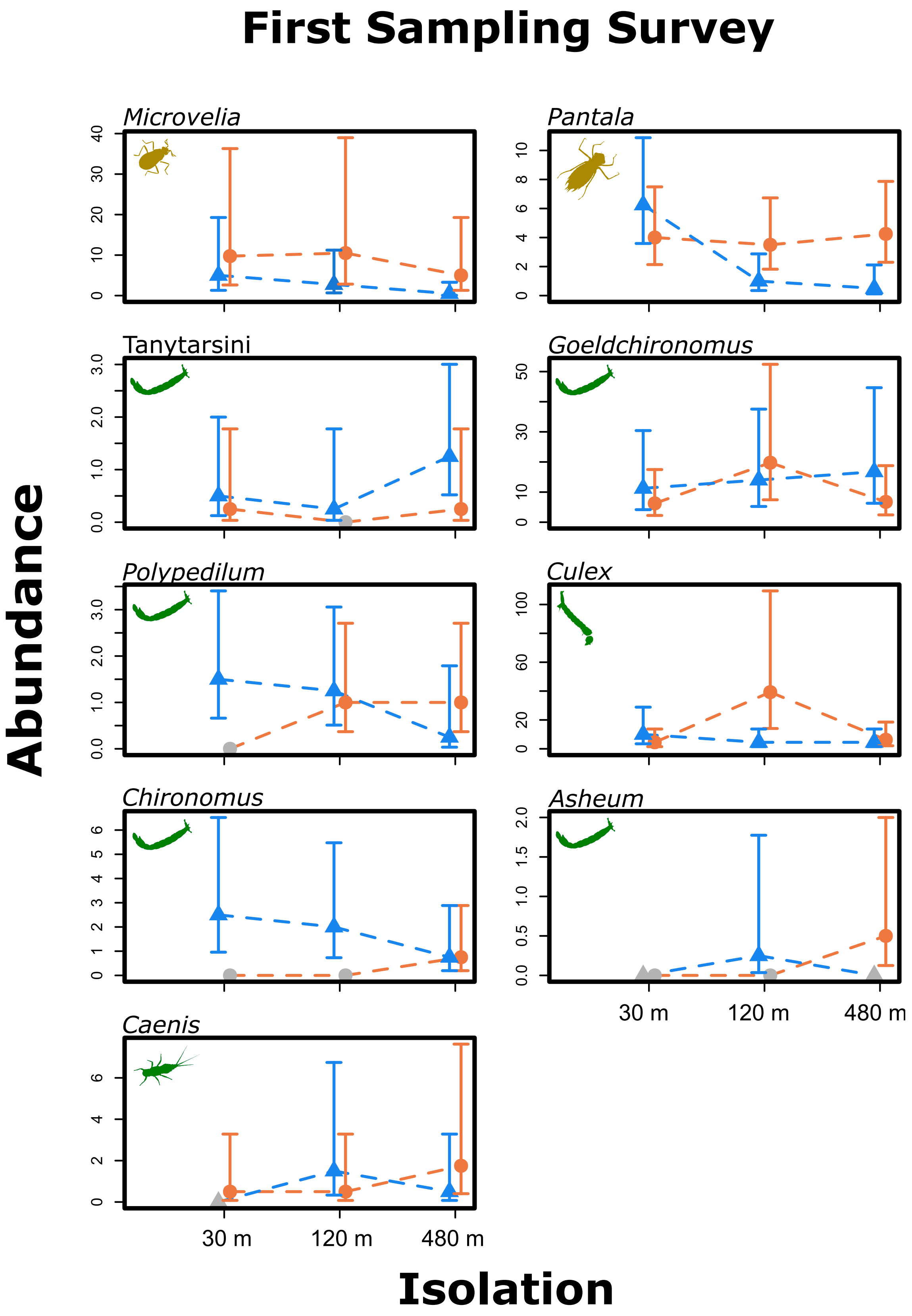

**Figure S9.3.** Effect of treatments on species abundance in the first sampling survey according to maximum likelihood estimates (MLE) of abundance and their 95% confidence interval for Model 8 in Table 1. Grey symbols represent absolute absence (zero abundance) of a taxon in a treatment. Blue triangles are MLEs for fish treatments and orange balls are MLEs for fishless treatments. The estimated differences are provided in Appendix 9.

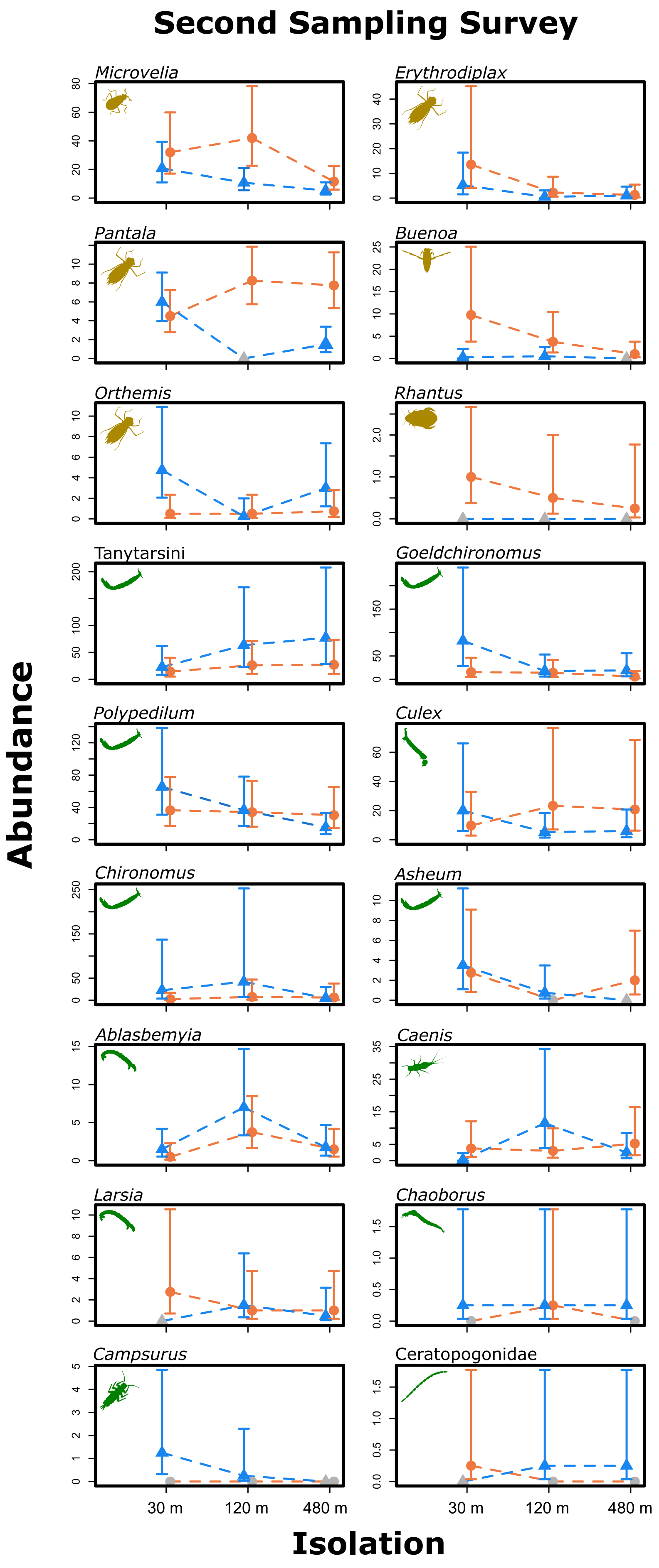

**Figure S9.4.** Effect of treatments on species abundance in the first sampling survey according to maximum likelihood estimates (MLE) of abundance and their 95% confidence interval for Model 11 in Table 1. Grey symbols represent absolute absence (zero abundance) of a taxon in a treatment. Blue triangles are MLEs for fish treatments and orange balls are MLEs for fishless treatments. The estimated differences are provided in Appendix 9.

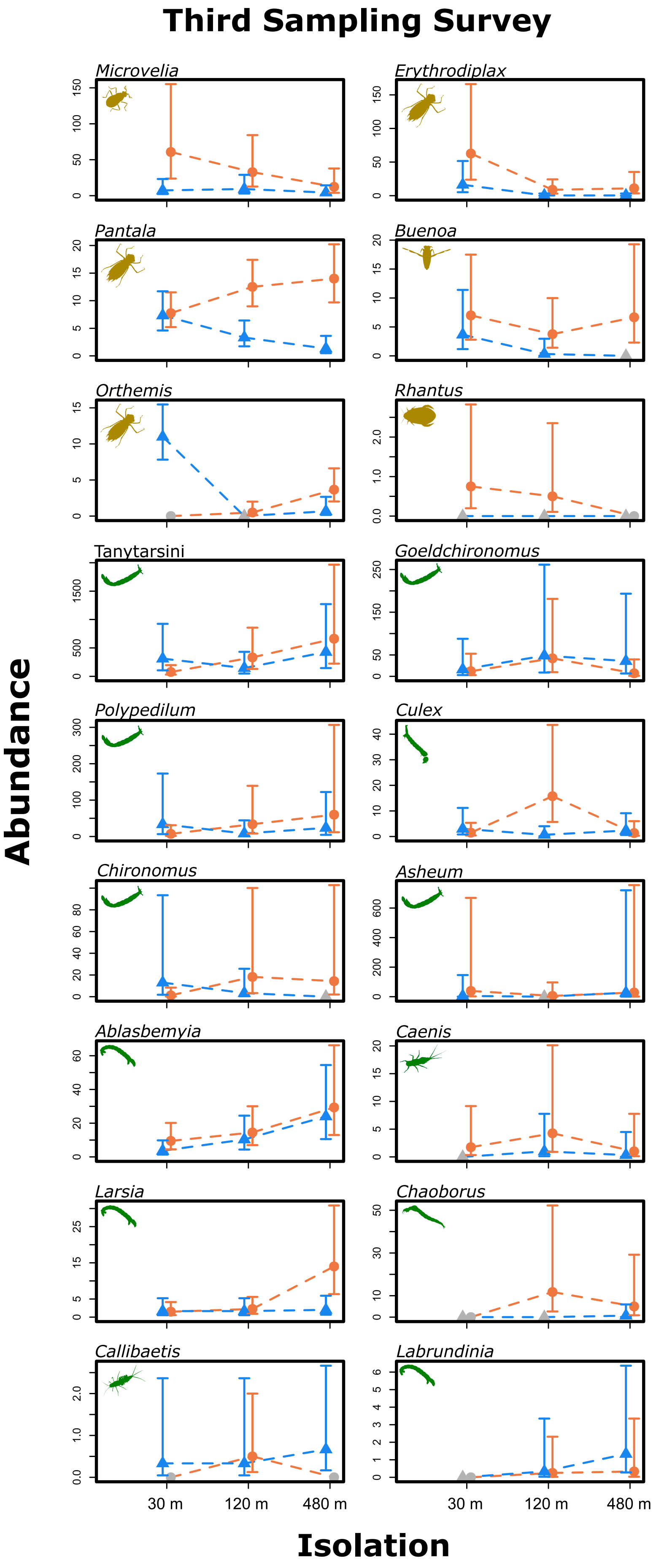

**Figure S9.5.** Effect of treatments on species abundance in the first sampling survey according to maximum likelihood estimates (MLE) of abundance and their 95% confidence interval for Model 15 in Table 1. Grey symbols represent absolute absence (zero abundance) of a taxon in a treatment. Blue triangles are MLEs for fish treatments and orange balls are MLEs for fishless treatments. The estimated differences are provided in Appendix 9.

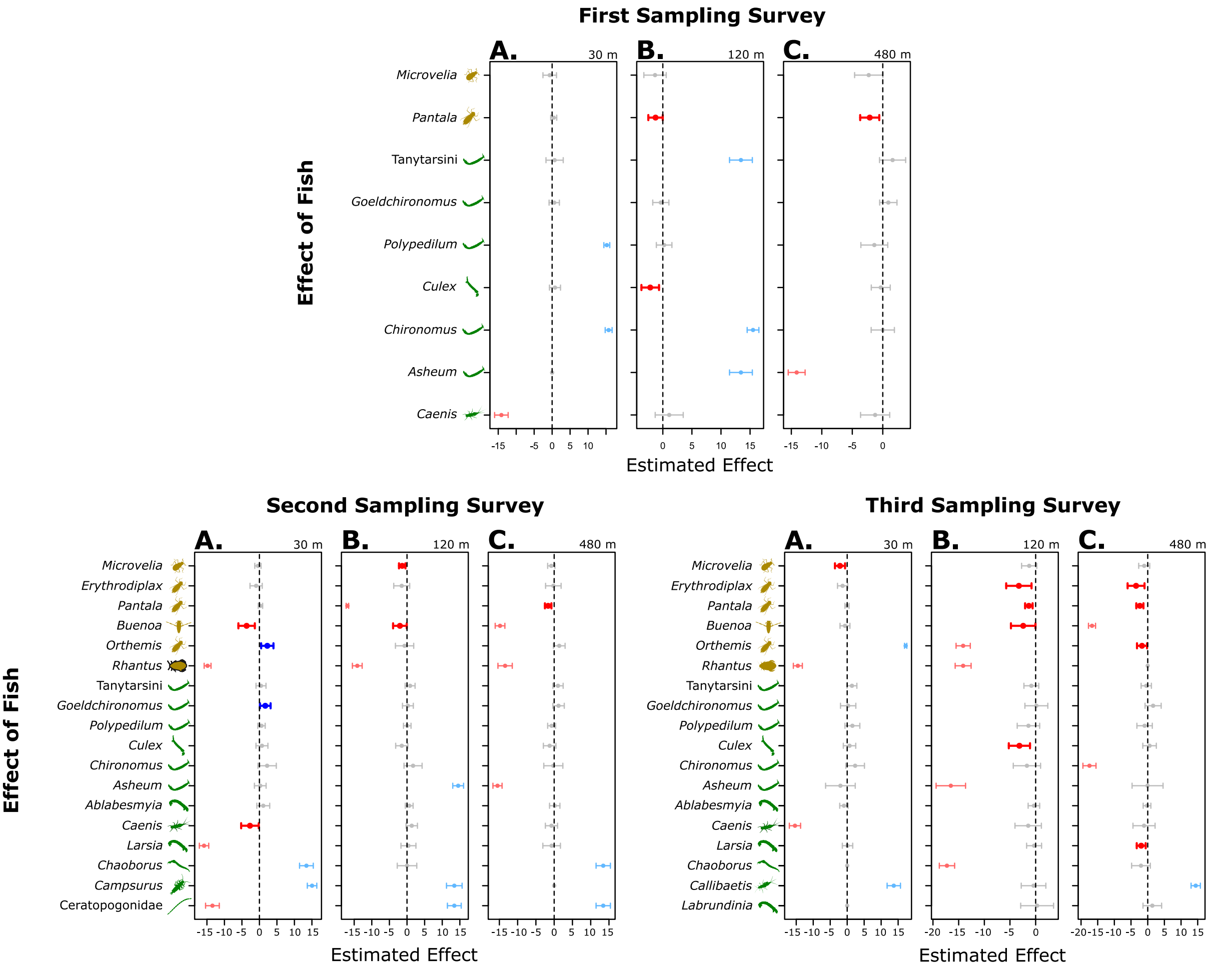

**Figure S9.6.** Confidence intervals for the effect of fish on abundance for each taxon, predatory insects (yellow), and herbivores and detritivores (green). A, B and C are effects for the first sampling survey, D, E and F are for the second and G, H and I are for the third. A, D and G are effects of the presence of fish in low isolation; B, E and H are effects of the presence of fish in moderate isolation; C, F and I are effects of the presence of fish in high isolation. See legend of Figure 1 for details.

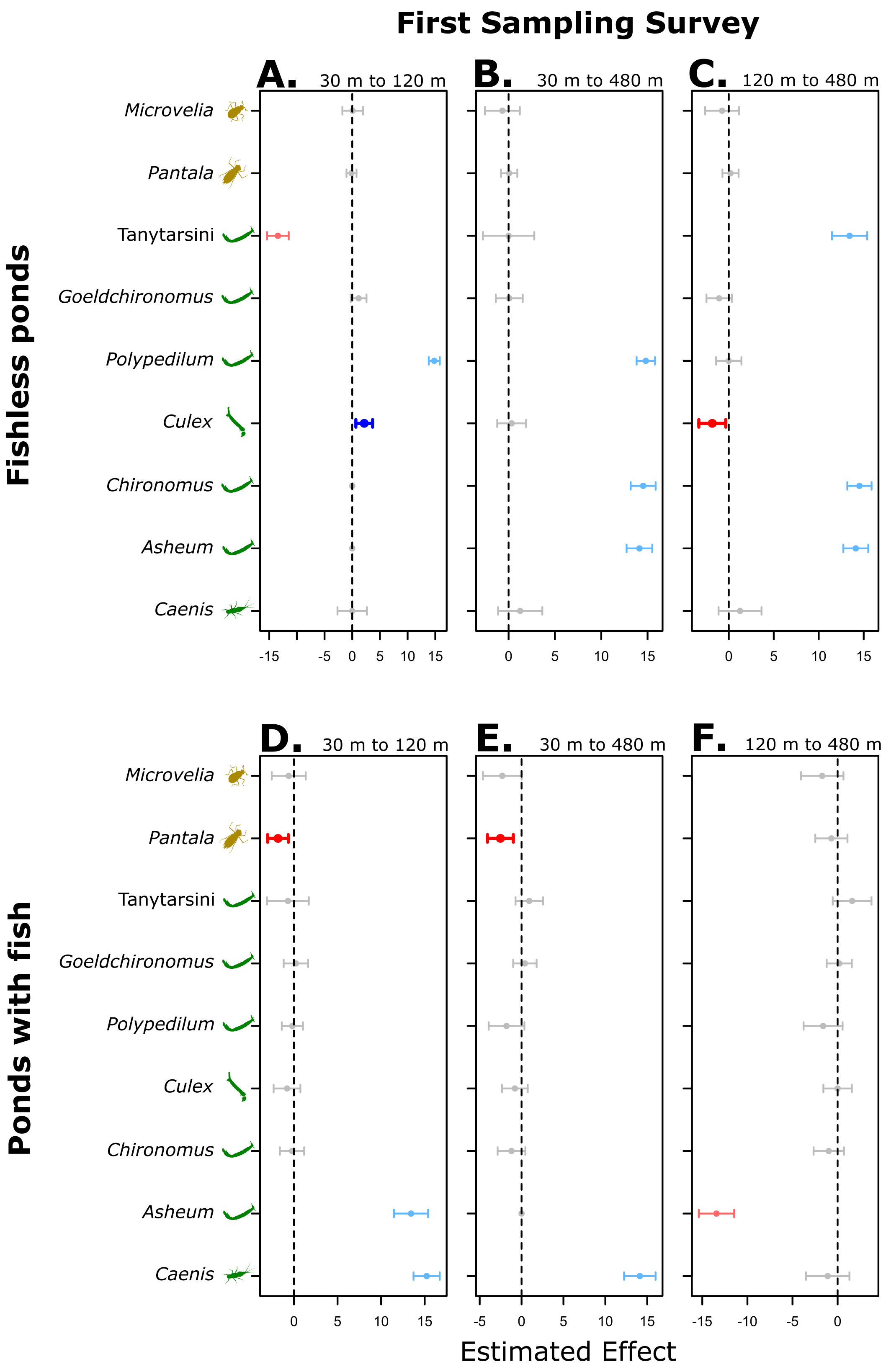

**Figure S9.7.** Confidence intervals for the effect of isolation on abundance for each taxon, predatory insects (yellow), and herbivores and detritivores (green), in the first sampling survey. A to C are effects for fishless ponds and D to F are for ponds with fish. A and D are effects of increasing isolation from 30 to 120 m; B and E are effects of increasing from 30 to 480 m. C and F are effects of increasing from 120 m to 480 m. See legend of Figure 1 for details.

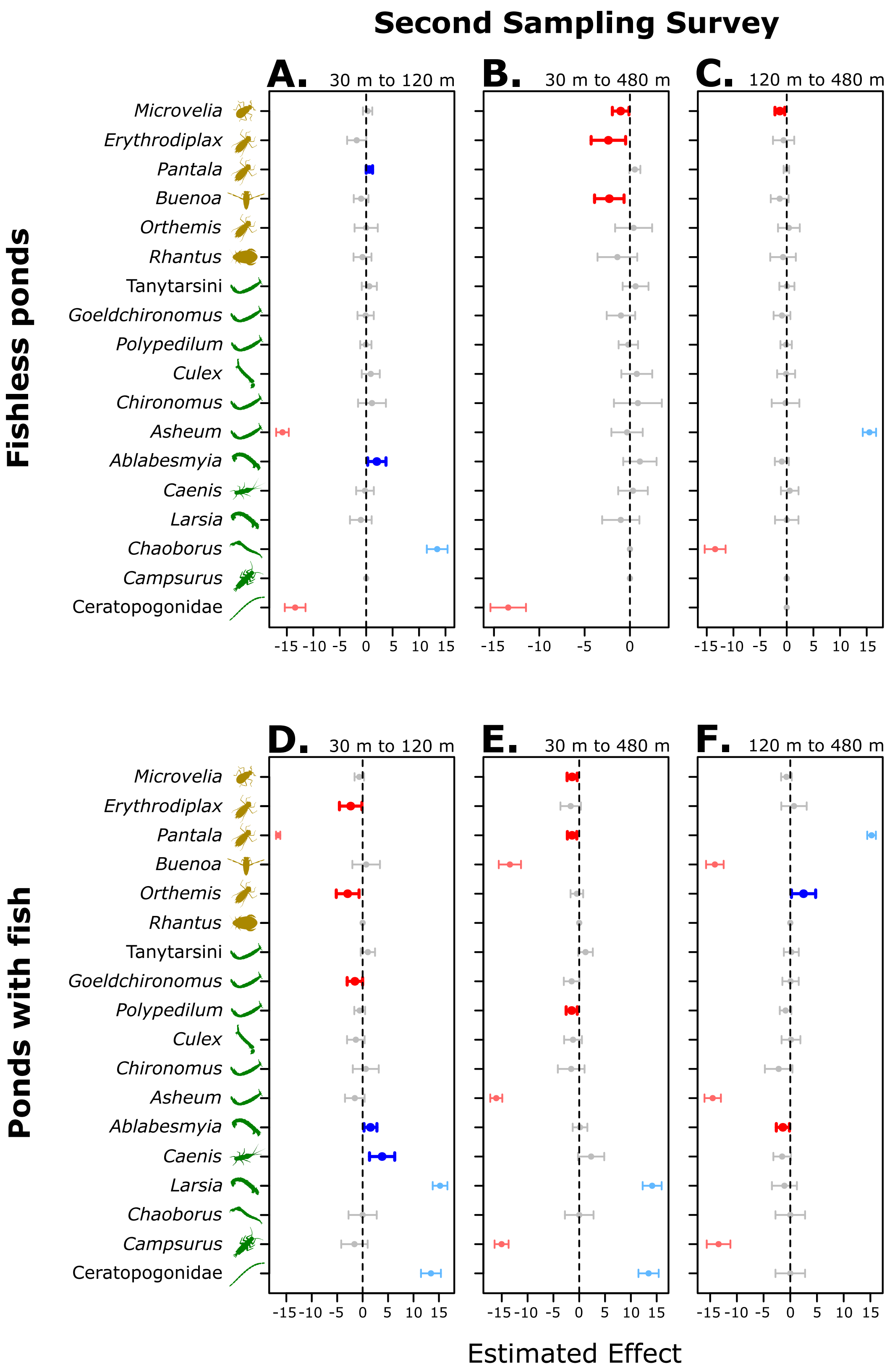

**Figure S9.8.** Confidence intervals for the effect of isolation on abundance for each taxon, predatory insects (yellow), and herbivores and detritivores (green) in the second sampling survey. A to C are effects for fishless ponds and D to F are for ponds with fish. A and D are effects of increasing isolation from 30 to 120 m; B and E are effects of increasing from 30 to 480 m. C and F are effects of increasing from 120 m to 480 m. See legend of Figure 1 for details.

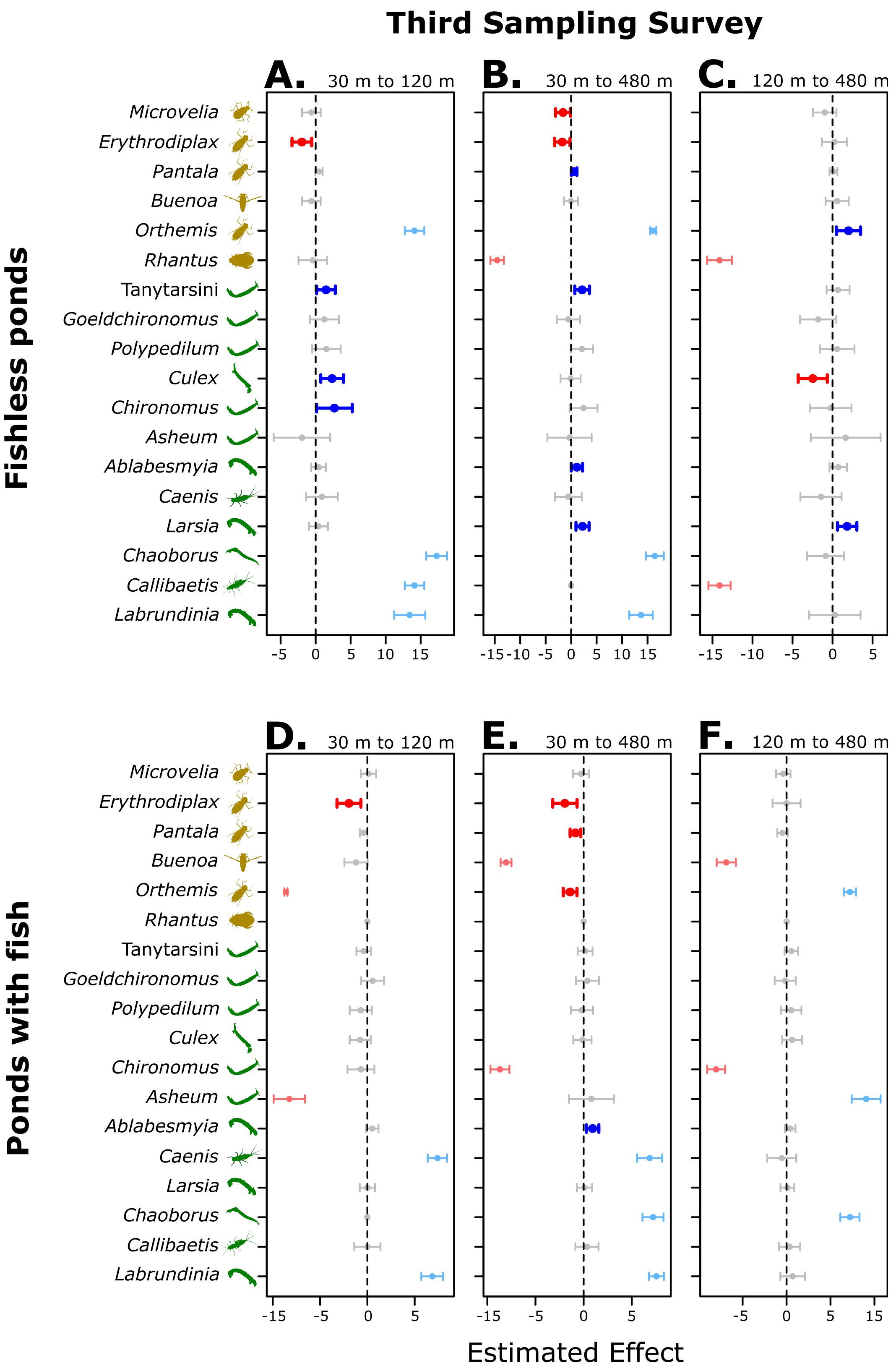

**Figure S9.9.** Confidence intervals for the effect of isolation on abundance for each taxon, predatory insects (yellow), and herbivores and detritivores (green) in the second sampling survey. A to C are effects for fishless ponds and D to F are for ponds with fish. A and D are effects of increasing isolation from 30 to 120 m; B and E are effects of increasing from 30 to 480 m. C and F are effects of increasing from 120 m to 480 m. See caption of Figure S8.2 for details.
